# Widening the bottleneck: Heterologous expression, purification, and characterization of the *Ktedonobacter racemifer* minimal type II polyketide synthase in *Escherichia coli*

**DOI:** 10.1101/2020.05.18.102780

**Authors:** Joshua G. Klein, Yang Wu, Bashkim Kokona, Louise K. Charkoudian

## Abstract

Enzyme assemblies such as type II polyketide synthases (PKSs) produce a wide array of bioactive secondary metabolites. While the molecules produced by type II PKSs have found remarkable success in the clinic, the biosynthetic prowess of these enzymes has been stymied by: 1) the inability to reconstitute the bioactivity of the minimal PKS enzymes *in vitro* and 2) limited exploration of type II PKSs from diverse phyla. Towards filling this unmet need, we expressed, purified, and characterized the ketosynthase chain length factor (KSCLF) and acyl carrier protein (ACP) from *Ktedonobacter racemifer*. Using *E. coli* as a heterologous host, we obtained soluble proteins in titers representing significant improvements over previous KSCLF heterologous expression efforts. Characterization of these enzymes reveals that *Kr*ACP has self-malonylating activity. Sedimentation velocity analytical ultracentrifugation (SV-AUC) analysis of *holo*-*Kr*ACP and *Kr*KSCLF indicates that these enzymes do not interact *in vitro*, suggesting that the acylated state of these proteins might play an important role in facilitating biosynthetically relevant interactions. Given the potential impact of obtaining soluble core type II PKS biosynthetic enzymes to enable *in vitro* characterization studies, these results lay important groundwork for optimizing the interaction between *Kr*KSCLF and *Kr*ACP and exploring the biosynthetic potential of other non-actinomycete type II PKSs.

## Introduction

Microorganisms produce secondary metabolites, small organic molecules that are not essential to the organism’s growth and reproduction but rather confer an evolutionary advantage. These molecules, often referred to as “natural products,” have been repurposed by humans as pharmaceutical agents and biological probes. Over 60% of all drugs approved between 1981 – 2014 originated from natural products and their synthetic derivatives, with notable examples including erythromycin (antibiotic), lovastatin (cholesterol-lowering), rapamycin (immunosuppressant), doxorubicin (anticancer) and vancomycin (antibiotic).^1–3^ Polyketides represent a remarkably successful class of natural products. Molecules in this class are manufactured by multi-enzyme pathways called polyketide synthases (PKSs), which are encoded by biosynthetic gene clusters. The chemical diversity programmed by polyketide biosynthetic gene clusters has been expanded through engineering efforts. For example, the diversity of molecules produced by type I PKSs (which are comprised of modular multidomain proteins and typically manufacture macrocyclic lactone polyketides) has been expanded via precursor-directed biosynthesis, substrate specificity engineering and combinatorial biosynthesis.^4,5^ Conversely, progress towards expanding the chemical diversity of type II PKSs (which are comprised of discrete proteins that act iteratively to manufacture polyaromatic polyketides) has been stymied by the difficulties in obtaining soluble biosynthetic proteins for structural and functional characterization. Nonetheless, type II PKSs remain a particularly attractive source of new chemical diversity because of their outstanding track record for producing pharmaceutically-relevant molecules.^1,6,7^

The core proteins involved in type II polyketide biosynthesis include the heterodimeric ketosynthase-chain length factor (KSCLF) and the acyl carrier protein (ACP) (**Fig 1**). These enzymes act collaboratively and iteratively to produce a nascent polyketide chain. The reactive β-keto chain is converted into a structurally complex molecule through the action of tailoring enzymes, including cyclases and ketoreductases, giving rise to the final branching, oxidation state, and cyclization pattern of the polyaromatic product. The remarkable chemical diversity observed in this class of molecules does not origin ate from monomer diversity, but rather from variations in chain length and tailoring reactions. In principle, molecules of novel structure and bioactivity can be created by altering the identity of the KSCLF (chain length), tailoring enzymes (oxidation state, cyclization pattern), and the presence of KSIII (substituents). However, there remain fundamental unanswered questions that limit our ability to engineer type II PKSs to make molecules of novel structure and function. For example, we only know pieces of the puzzle as to how type II PKSs guide the efficient vectorial channeling of polyketide intermediates to the appropriate enzymes, which directs the structure of products.^8^ In addition, unnatural protein-protein and protein-substrate interactions are often insufficient to generate catalytically efficient synthases.^9,10^ Engineering efforts are further hampered by the large and growing deficit in functionally annotated enzymes.^5^

**Fig 1.**
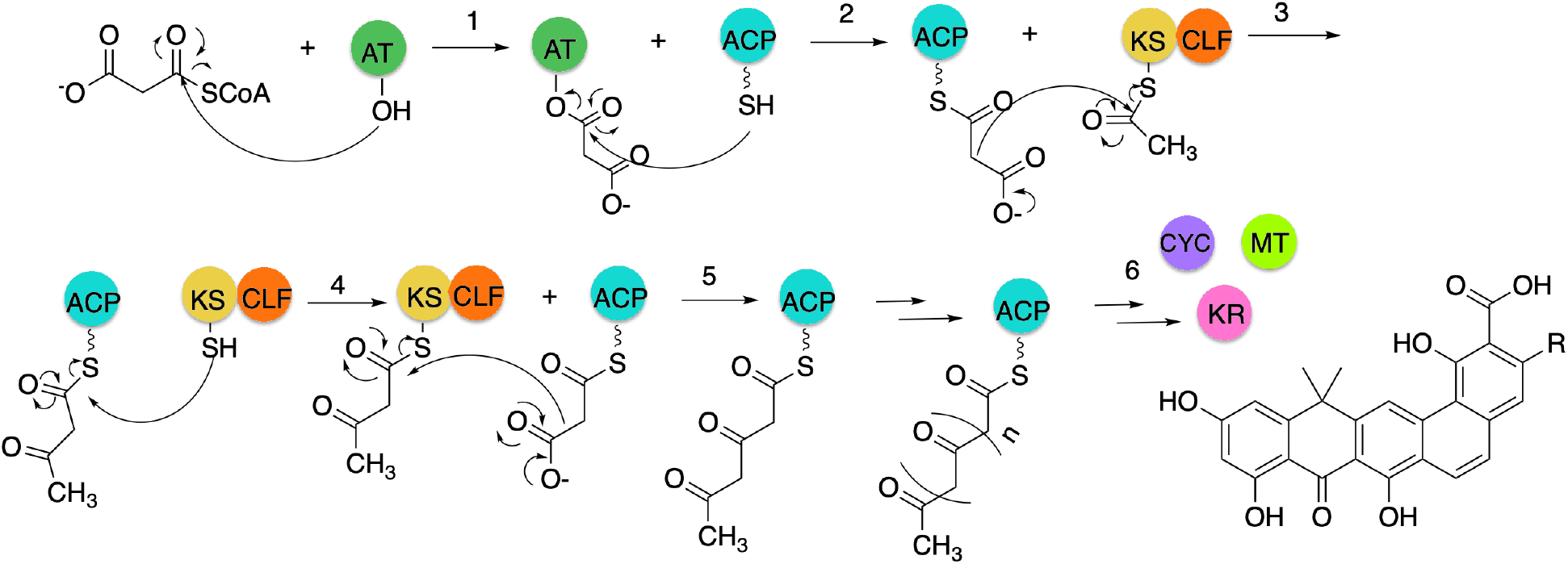
Overview of the role of the KSCLF and ACP in type II polyketide biosynthesis. 1. If necessary, an acyl transferase (AT) reacts with malonyl-coenzyme A (malonyl-CoA) via a nucleophilic acyl substitution reaction to malonylate the AT. 2. The terminal thiol group on the ACP phosphopantetheine arm (represented as a squiggly line) reacts via a nucleophilic acyl substitution reaction to malonylate the ACP. If the ACP is “self-malonylating”, the terminal thiol can directly load the malonyl-CoA. 3. Malonylated-ACP reacts with the acylated active thiol on the KSCLF via a decarboxylative Claisen-like condensation reaction to initiate the formation of the polyketide chain on the ACP. 4. The polyketide chain is transferred to the KSCLF via an acyl substitution reaction. 5. The β-keto-bound KSCLF reacts with malonylated ACP via a decarboxylative Claisen-like condensation to further elongate the polyketide chain; this process repeats until the polyketide chain is elongated to the programmed chain length, which in part is directed by the steric cavity of the KSCLF.^11^ 6. Tailoring enzymes such as cyclases (CYC), methyltransferases (MT), and ketoreductases (KR) act on the elongated polyketide chain to form the final natural product.

In an effort to study the evolution of biosynthetic genes on a cluster-wide scale, we previously created a catalog of genes that putatively encode for type II PKS enzymes.^11^ This work, which was led by Hillenmeyer and conducted in collaboration with our research team, unveiled that the enormous structural and biological diversity of type II polyketide antibiotics originated from primary metabolic fatty acid synthases (FASs).^11^ In our studies, we noted that at the time all characterized polyketide gene clusters (sequenced gene clusters whose secondary metabolite products had been structurally characterized as of 2015) were from actinobacteria. We also revealed that the divergence of type II PKSs from FASs predated major speciation events, suggesting that type II PKSs diverged from FASs well before the actinobacterial phylum had formed. This led us to recognize the clade of uncharacterized, ancient type II PKS biosynthetic gene clusters from non-actinobacterial species as unexplored space ripe for the mining of molecules of novel structure and function, as well as enzymes with novel biocatalytic capabilities.

We were particularly excited about the possibility of characterizing KSCLFs from non-actinobacteria because this heterodimer plays an essential role in the manufacturing of type II PKSs but has long evaded robust expression and purification methods. The challenges posed by the lack of a robust heterologous expression platform for the type II PKSs has hampered progress towards understanding and engineering these powerful biosynthetic systems.^12^ We hypothesized that the ancient type II PKS genes harbored in non-actinobacterial species might be amenable to expression in *E. coli* and initiated work in this arena. Recent groundbreaking work by Cummings and co-workers confirmed that indeed these non-actinobacterial type II PKS gene clusters are ripe for bioprospecting.^13^ The researchers showed that type II PKS enzymes from non-actinomycetes can be expressed in *E. coli* and that these enzymes can effectively biosynthesize type II polyketides *in vitro*. Moreover, the authors showed that various tailoring enzymes can be substituted into the reconstituted type II PKS assembly to alter the final type II polyketide. These findings provide an exciting platform for the future synthetic biology efforts and also inspired us to pivot our research focus towards gaining a molecular-level understanding of the structure and function of core type II PKS enzymes.

Herein, we present initial results on the characterization of type II PKS biosynthetic enzymes from the non-actinobacterium *Ktedonobacter racemifer* DSM 44963. *K. racemifer* is a gram-positive bacterium belonging to the Chloroflexi phylum, isolated from Italian forest soil.^14^ It harbors a putative type II PKS biosynthetic gene cluster that belongs to the ancient resistomycin clade;^11^ we viewed its evolutionary proximity to the *E. coli* FAS as a harbinger that the core enzymes might be amenable to expression in *E. coli*. We report optimized expression conditions for the *K. racemifer* minimal PKS components, *Kr*ACP and *Kr*KSCLF, in *E. coli*. Successful expression of *Kr*ACP and *Kr*KSCLF enabled their biophysical characterization and set the stage for future investigations into their biocatalytic potential.

## Results

### Expression and purification of *K. racemifer* minimal PKS enzymes

The *K. racemifer* minimal PKS enzymes, *Kr*ACP and *Kr*KSCLF were successfully expressed and purified from the *E. coli* BAP1^15^ and BL21 competent cell lines, respectively (**Fig 2**). Titers of 16 mg/L for *Kr*ACP and 3 mg/L for *Kr*KSCLF were obtained from purification. Expression of *Kr*ACP in the BAP1 cell line, which harbors the broad substrate phosphopantetheinyl transferase from *B. subtilis* Sfp, afforded ~50 % of the ACP with phosphopantetheine (Ppant) arm installed (“*holo*-*Kr*ACP”) and ~50 % *apo*-*Kr*ACP. (**Fig S1, S2**). Species identities of *holo-Kr*ACP and *apo-Kr*ACP upon analysis by high-performance liquid chromatography (HPLC) are supported by literature characterization of other ACPs and by spiking experiments performed using *E. coli holo*-AcpP and *apo*-AcpP (**Fig S3**).^16,17^ Presence of *holo*-*Kr*ACP was also confirmed by sodium dodecyl sulfate-polyacrylamide (SDS-PAGE) gel electrophoresis, as the ~27 kDa band corresponding to a disulfide bond-linked *holo*-*Kr*ACP dimer is observed under non-reducing conditions and not in the presence of 5% v/v β-mercaptoethanol (BME; **Fig 2A**).

**Fig 2.**
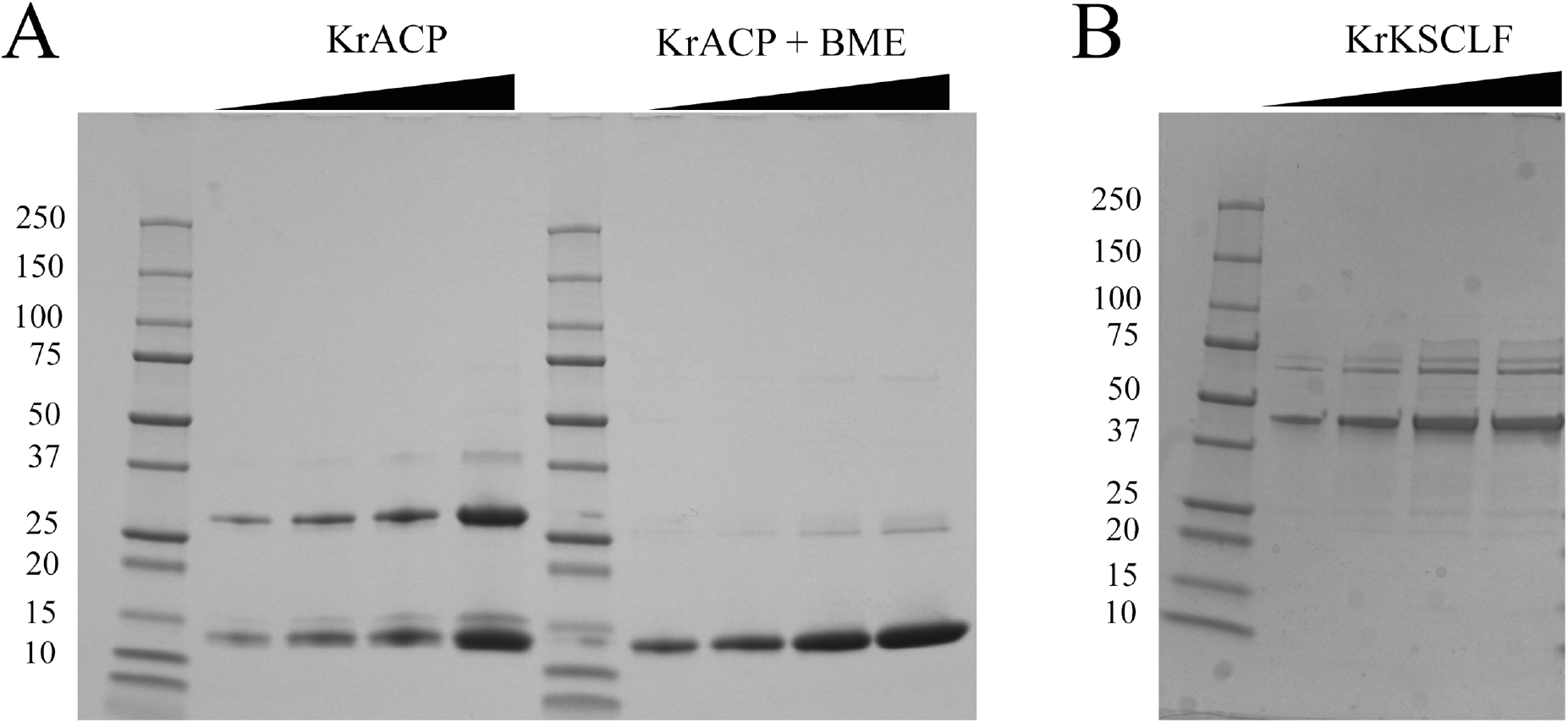
Characterization of *K. racemifer* minimal PKS enzymes by SDS-PAGE. A. *Kr*ACP concentration gradients run both without and with the presence of 5% v/v BME. Lanes: 1. Protein standards ladder; 2. *Kr*ACP (31 μM, 5 μL) run under non-reducing conditions; 3. *Kr*ACP (31 μM, 10 μL) run under non-reducing conditions; 4. *Kr*ACP (31 μM, 15 μL) run under non-reducing conditions; 5. *Kr*ACP (31 μM, 20 μL) run under non-reducing conditions; 6. Protein standards ladder; 7. *Kr*ACP (31 μM, 5 μL) run under reducing conditions; 8. *Kr*ACP (31 μM, 10 μL) run under reducing conditions; 9. *Kr*ACP (31 μM, 15 μL) run under reducing conditions; 10. *Kr*ACP (31 μM, 20 μL) run under reducing conditions. The disappearance of the ~27 kDa band upon the addition of reducing agent indicates that this band represents the *holo*-*Kr*ACP disulfide dimer. B. *Kr*KSCLF purified under reducing conditions. 1. Protein standards ladder; 2. *Kr*KSCLF (5 μL, 2.6 μM); 3. *Kr*KSCLF (10 μL, 2.6 μM); 4. *Kr*KSCLF (15 μL, 2.6 μM); 5. *Kr*KSCLF (20 μL, 2.6 μM). The ~47 kDa band was excised and confirmed to contain the *Kr*KS and *Kr*CLF monomers via tandem proteolysis mass spectrometry (Figures S4). Purification under reducing conditions was necessary to avoid aggregation and purification of the nickel column elute via anion exchange was required to remove ~25 kDa *E. coli* protein contaminants (see Results and Supporting Information for details).

Far UV-Circular Dichroism spectroscopy (CD) was used to determine the secondary structure of *Kr*ACP (**Fig 3A**). Similar to other ACPs studied in our laboratory, *Kr*ACP in solution is mostly helical with two negative peaks at 222 nm and 208 nm and a positive peak at 192 nm (**Fig 3A**).^18^ Pre- and post-thermal denaturation spectra showed that unfolding is reversible with greater than 80% recovery in signal following denaturation (**Fig 3A**). Thermal unfolding appeared to be a cooperative two-state process without observable intermediates with a midpoint T_m_ = 66.7 ± 1.8 °C, enthalpy of unfolding ΔH_m_ = 118.7 ± 13.4 kJ mol^−1^, and change in entropy ΔS_m_ = 1.8 ± 0.2 kJ (K^−1^ mol^−1^) (**Fig 3A inset**). Sedimentation velocity experiments with the analytical ultracentrifuge (SV-AUC) indicates that *Kr*ACP sediments mostly as a monomer with a molecular weight of ~12 kDa (theoretical molecular weight of *holo Kr*ACP is 12.2 kDa; **Fig 3B**). The sedimentation of *Kr*ACP in primarily monomeric form is consistent with what has been observed for other type II PKS ACPs.^18^

**Fig 3.**
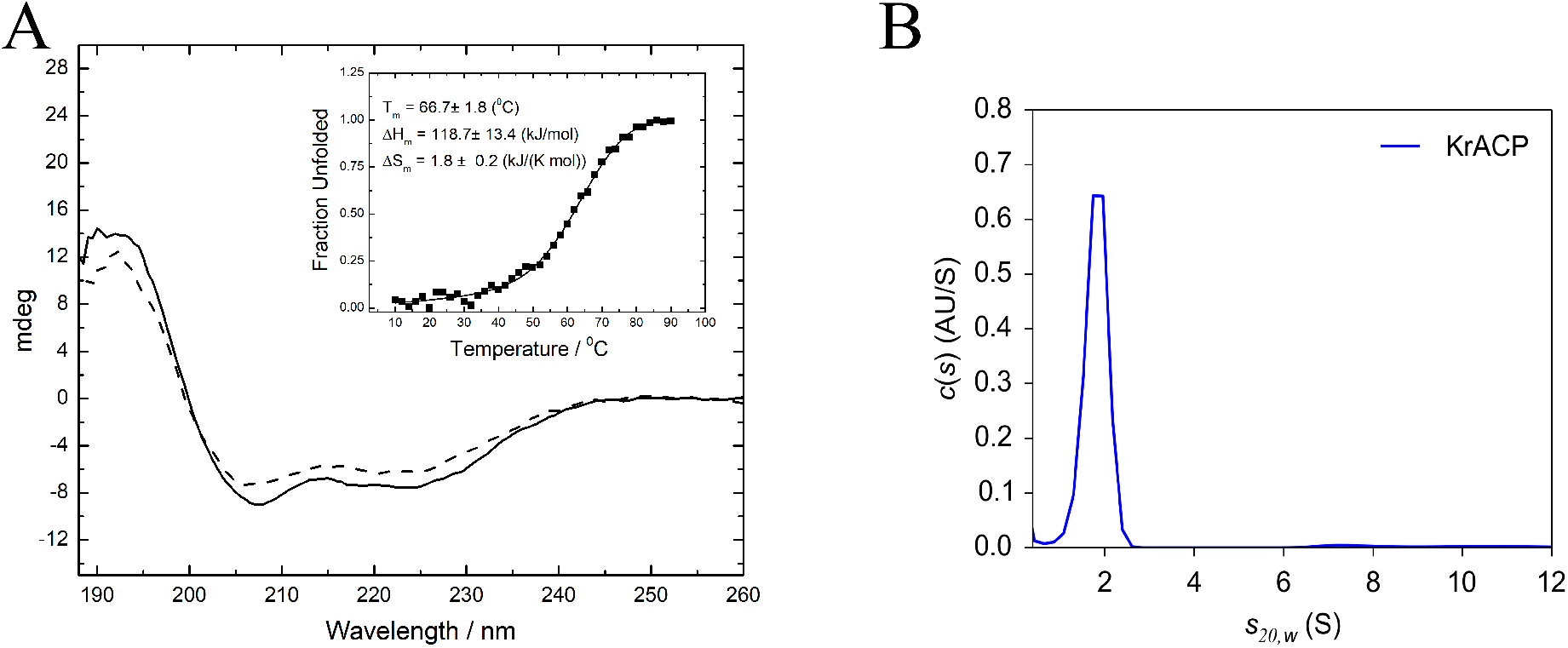
Characterization of *Kr*ACP. A. Circular dichroism (CD) spectra and melting temperature experiments of *Kr*ACP. The CD spectrum of purified *Kr*ACP is given by the solid line, while the CD spectrum of *Kr*ACP after melting temperature experiments is given by the dashed line. Melting temperature curve measured at 222 nm and subsequent extracted thermodynamic parameters are given. The similarity in pre- and post-heating CD spectra indicate that *Kr*ACP folds to regain nearly all of its secondary structure after heating to 90 °C. B. SV-AUC profile of *Kr*ACP under reducing conditions. *Kr*ACP shows as a single species in SV-AUC with a molecular weight of ~12 kDa, indicating that the protein is monomer in solution and that any disulfide bonds between *holo*-*Kr*ACP monomers are reduced through the presence of dithiothreitol (1 mM DTT) in the buffer (50 mM sodium phosphate, pH 7.6, 150 mM NaCl).

Initial *Kr*KSCLF purification attempts under non-reducing conditions led to protein aggregation. The addition of BME to the samples reduced aggregation, suggesting that aggregate formation was at least partially caused by nonspecific cysteine interactions between *Kr*KSCLF monomers. As such, the expression protocol was optimized to include the addition of reducing agent in all purification and storage buffers. SDS-PAGE gel electrophoresis of the reducing buffer-purified *Kr*KSCLF revealed three distinct bands at ~ 85 kDa, 47 kDa, and 25 kDa (**Fig S4).** The highest molecular weight (MW) band, corresponding to a mass of approximately 85 kDa, was attributed to the *Kr*KSCLF heterodimer. The ~47 kDa band and ~25 kDa bands were excised for analysis by tandem-proteolysis mass spectrometry. The ~47 kDa band was determined to be composed of both the *Kr*KS and *Kr*CLF monomers (**Fig S5**). The ~25 kDa band was partially composed of the *Kr*KS and *Kr*CLF monomers, but primarily composed of three proteins native to *E. coli*: FKBP-type peptidyl prolytl cis-trans isomerase SlyD (NCBI accession number P0A9K9), hydroxyethylthiazole kinase ThiM (NCBI accession number P76423), and triosephosphate isomerase TpiA (NCBI accession number P0A858) **Fig S6**). This ~25 kDa species was also observed in the SV-AUC profile of *Kr*KSCLF sample prior to further purification of the sample under reducing conditions by size exclusion chromatography (SEC; **Figs S4 and S7)**. Concentration dependence SV-AUC revealed that the ratio of *Kr*KSCLF to 25 kDa proteins remained unchanged regardless of the concentration of sample loaded. These findings indicate that the identified *E. coli* proteins co-purified with *Kr*KSCLF and co-precipitated with the *Kr*KSCLF heterodimer to the extent that we were not able to separate via SEC (in absence of salt and reducing agent). Inclusion of 150 mM NaCl and 1 mM DTT in the SEC buffer made it possible to purify the *Kr*KSCLF proteins away from the *E. coli* contaminants. Subsequent analysis of the purified *Kr*KSCLF heterodimer by CD indicates that the secondary structure is comprised of both β-sheets and ɑ-helices (**Fig 4A**). Whereas the *Kr*KS and *Kr*CLF run as a monomer-dimer equilibrium via SDS PAGE (**Fig 2**), the proteins sediment as the *Kr*KSCLF heterodimer in solution (**Fig 4B**).

**Fig 4.**
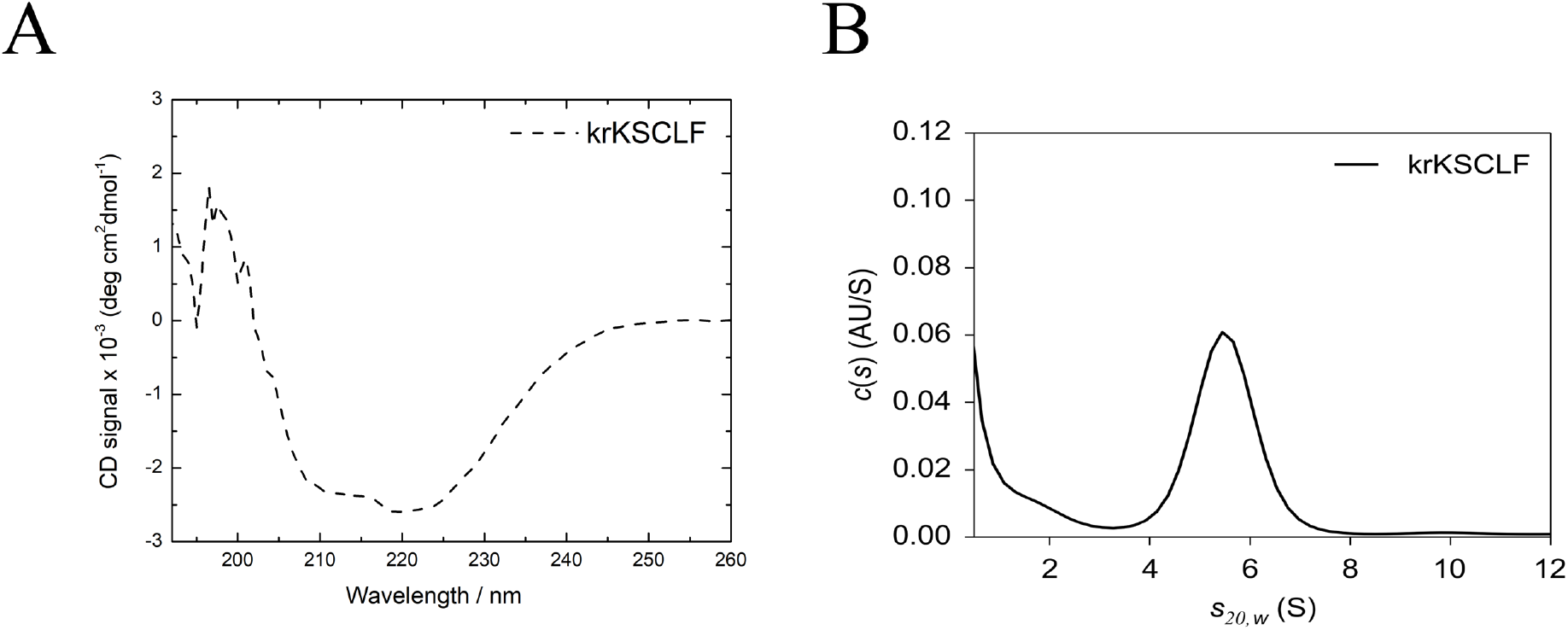
Characterization of *Kr*KSCLF. A. The CD spectrum of purified *Kr*KSCLF indicates that the heterodimer is comprised of both β-sheets and ɑ-helices. B. SV-AUC analysis of purified *Kr*KSCLF. The SV-AUC profile of purified *Kr*KSCLF reveals the presence of a major species of ~98.7 kDa, which is roughly consistent with the molecular weight of the *Kr*KSCLF heterodimer (theoretical molecular weight of 92.6 kDa)

### Malonylation of *Kr*ACP

To initiate biosynthesis, the Ppant arm of *holo*-ACP must be primed with an initial malonyl extender unit (**Fig 1**). In many well-studied type II PKSs, this *holo*-ACP malonylation is performed by an AT or malonyl transferase, such as *E. coli* FabD.^19,20^ However, some ACPs have previously been reported malonylate without the presence of a malonyl transferase.^21^ The ability of the *Kr*ACP to undergo this so-called self-malonylation was assessed by characterizing the products of *Kr*ACP and malonyl-CoA in the presence and absence of FabD. The reaction solutions were purified by SEC, and the fraction corresponding to *Kr*ACP were collected for analysis by liquid chromatography-mass spectrometry (LCMS) (**Fig S8**). LCMS samples were analyzed using a modified Ppant ejection assay.^22^ Both reaction solutions analyzed by this Ppant ejection assay revealed species with *m/z* values of 347.1 Da, corresponding to the malonylated, cyclized Ppant arm (**Fig 5**). The presence of malonylated *Kr*ACP in both samples was additionally confirmed by the deconvolution of higher *m/z* species corresponding to the molecular weight of malonylated *Kr*ACP (**Fig S9**). Taken together, these data suggest that *Kr*ACP can load malonyl CoA without the presence of a transferase partner. Whether or not this self-malonylation mechanism is physiologically relevant has yet to be determined.

**Fig 5.**
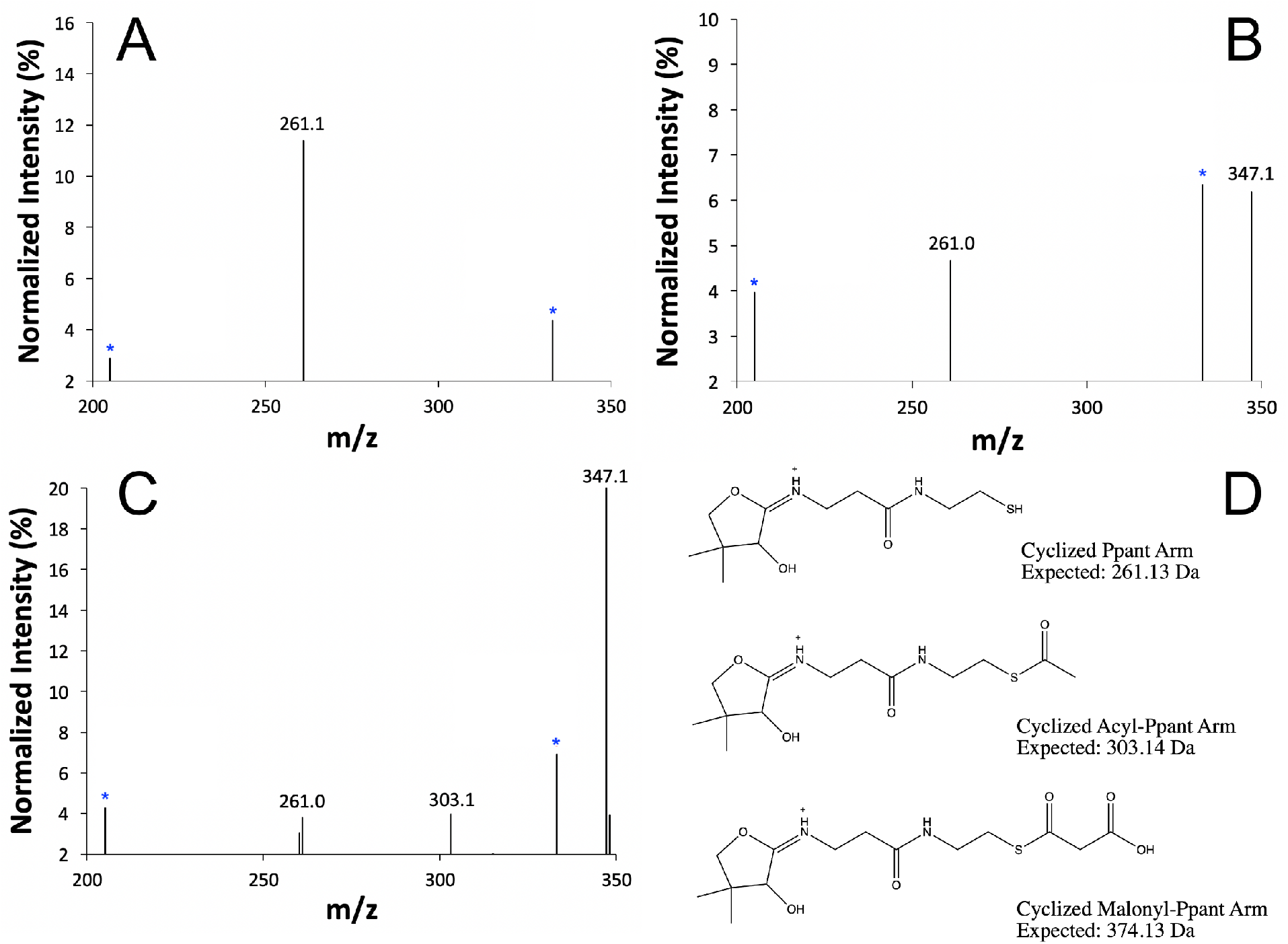
*Kr*ACP can load malonyl CoA in absence of an acyl transferase. A. Ppant ejection of *Kr*ACP shows the presence of *holo*-*Kr*ACP, represented by the 261.1 Da species corresponding to the cyclized Ppant arm. B. Ppant ejection of *Kr*ACP incubated with malonyl-CoA shows the appearance of a 347.1 Da species corresponding to the cyclized, malonylated Ppant arm, indicating that *holo*-*Kr*ACP can load malonyl-CoA in the absence of a transferase. C. Ppant ejection of *Kr*ACP with malonyl-CoA and FabD shows the presence of both 261.1 Da and 347.1 Da peaks, with the addition of a 303.1 Da peak corresponding to a cyclized acyl-Ppant arm, which forms upon the decarboxylation of malonyl-*Kr*ACP. D. Expected species in mass spectra resulting from cyclization of *Kr*ACP Ppant arm. Species marked by a blue asterisk were present in all spectra, including buffer, and likely represent background noise apparent at high electron multiplier voltage.

### Lack of observed *holo*-*Kr*ACP/*Kr*KSCLF interaction *in vitro*

To investigate the interactions between *holo*-*Kr*ACP and *Kr*KSCLF *in vitro*, both enzymes were studied by SV-AUC under reducing conditions. As noted above, *Kr*ACP sediments as a single species corresponding to the *Kr*ACP monomer (**Fig 3B**) and the *Kr*KSCLF as a heterodimer (**Fig 4B**). A combined sample consisting of both *Kr*ACP (100 μM) and *Kr*KSCLF (4 μM) showed no significant shift in the higher MW peak corresponding to the *Kr*KSCLF heterodimer (**Fig 6**), suggesting a lack of interaction between *Kr*KSCLF and *holo*-*Kr*ACP *in vitro.*

**Fig 6.**
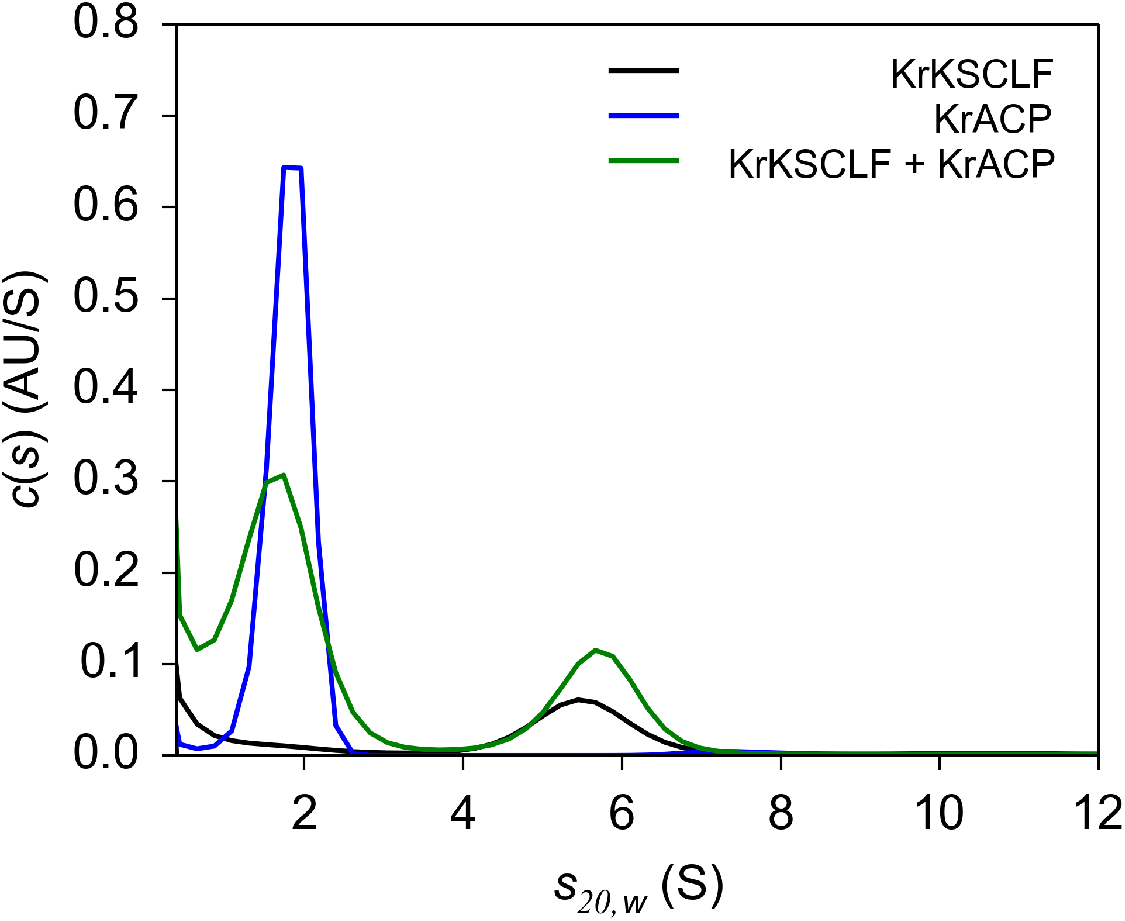
SV-AUC analysis of *K. racemifer* minimal PKS enzymes suggests a lack of interaction between *Kr*KSCLF and *Kr*ACP *in vitro*. SV-AUC analysis of minimal PKS enzymes showed no significant shift in the ~98.7 kDa peak corresponding to *Kr*KSCLF upon the addition of *Kr*ACP, indicating a lack of interaction between the enzymes under the investigated conditions. Conditions: 2.9 μM *Kr*KSCLF, 140 μM *Kr*ACP, and 4 μM *Kr*KSCLF combined with 110 μM *Kr*ACP. Samples were in 50 mM sodium phosphate, pH 7.2, 150 mM NaCl, and 1 mM DTT.

The lack of observed interactions between *holo*-*Kr*ACP and *Kr*KSCLF led us to further explore *Kr*KSCLF via protein modeling. *Kr*KSCLF was threaded onto the solved actKSCLF from *S. coelicolor*, which is currently the only polyaromatic type II PKS KSCLF available in the Protein Data Bank (**Fig 7**).^23^ The 17 Å amphipathic polyketide tunnel between Phe116 and Cys169 in actKSCLF has been shown to play a vital role in the elongation of the polyketide chain, with widening of the tunnel by site-directed mutagenesis corresponding to an increase in polyketide chain length.^23^ The threaded structure of *Kr*KSCLF shows the presence of a similar polyketide tunnel between residues Leu125 and Cys172; however, the width of this proposed *Kr*KSCLF amphipathic polyketide tunnel measures 22.5 Å. It is currently unclear if this predicted difference in tunnel shape affects how the *Kr*KSCLF binds to its cognate *Kr*ACP. A recent structure of the highly reducing type II PKS KSCLF Iga11-Iga12 enables us to draw some additional comparisons (**Fig S10**). First, both the polyketide tunnel size and charge of residues lining the cavity of Iga11-Iga12 were implicated in directing the length and structure of the manufactured product.^24^ In the Iga system, β-ketoacylation-ACP elongation is inhibited by Asp113, which does not appear to be conserved *Kr*KSCLF nor in other aromatic polyketide-synthesizing type II PKSs.^23,24^ However, the hydrogen bonding amino acids lining the IgaPKS KSCLF tunnel that stabilize the ACP Ppant arm (Thr 303 and Thr 305) appear to be conserved in *Kr*KSCLF. Second, the catalytic triad (Cys170-His301-His336) of Iga11 shifted upon acylation of and ACP binding.^24^ These comparisons support the possibility that *Kr*KSCLF can accommodate the biosynthesis of a polyaromatic polyketide, but that the acylation state of the *Kr*KSCLF and/or *Kr*ACP could play a pivotal role in ACP-KSCLF binding. Notably, the importance of enzyme acylation state in carrier protein interactions has been previously observed in other biosynthetic systems, including type I PKSs^25^ and non-ribosomal peptide synthetase (NRPSs).^26^

**Fig 7.**
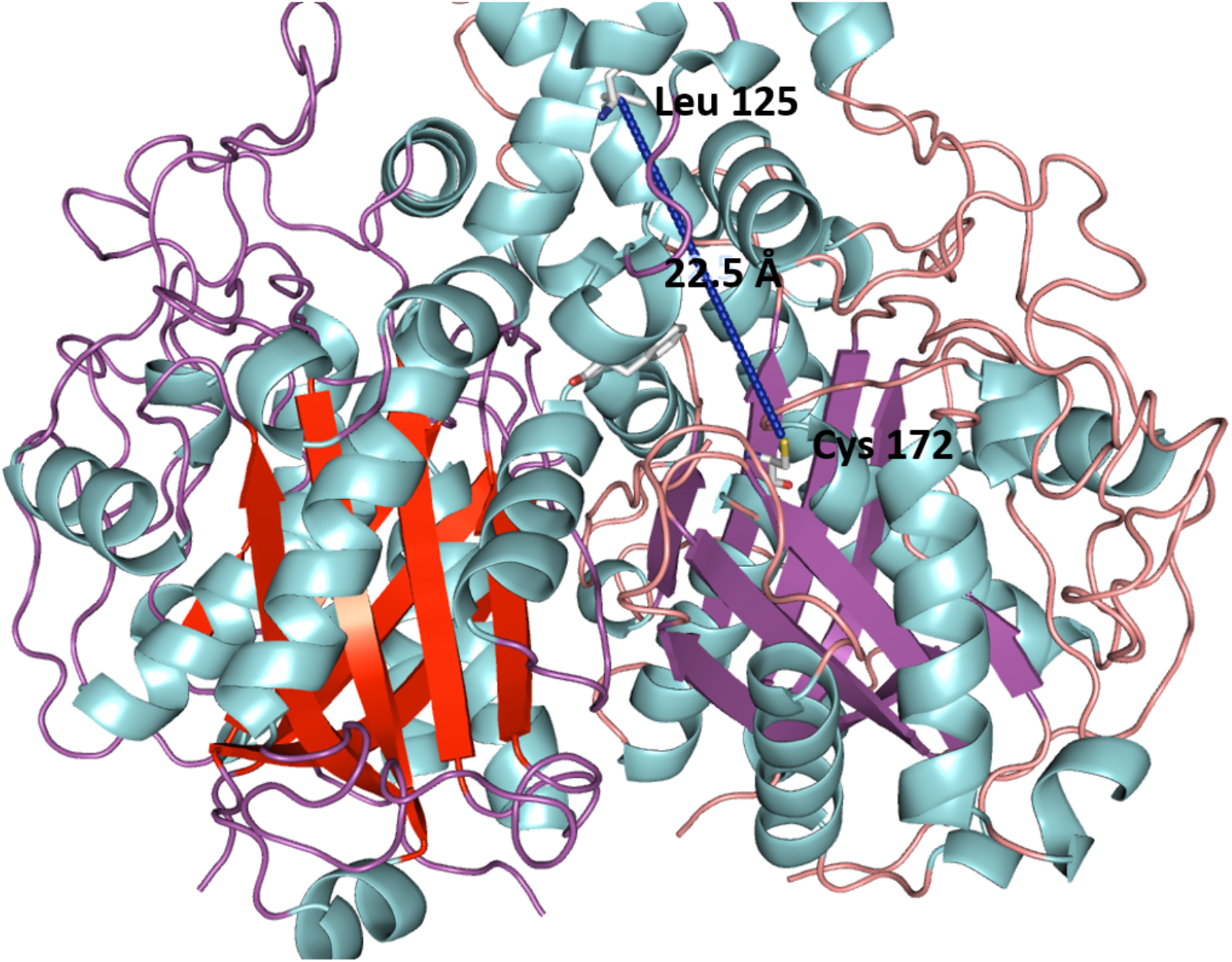
The *Kr*KSCLF sequence threaded onto the solved actKSCLF crystal structure (PDB 1TQY).^23^. The threaded *Kr*KSCLF structure reveals the heterodimer might harbor a similar polyketide tunnel to that of the previously studied in actKSCLF. However, the length of the proposed *Kr*KSCLF polyketide tunnel is 22.5 Å in comparison to the 17 Å polyketide tunnel of actKSCLF.23 It is possible that the length of the *Kr*KSCLF polyketide tunnel and/or the acylation state of the *Kr*KSCLF affects its ability to interact with the *holo*-*Kr*ACP.

## Discussion

The ability to heterologously express KSCLF enzymes in *E. coli* represents a longstanding bottleneck in the chemical engineering of type II PKSs. Recent developments have shown the ability of yeast and *E. coli* to serve as heterologous hosts for total type II PKS gene cluster reconstitution; however, the biochemical properties of the heterologously expressed minimal PKS enzymes from these systems have not been investigated.^13,27,28^ The heterologous expression of the *K. racemifer* minimal PKS enzymes *Kr*ACP and *Kr*KSCLF in *E. coli* is a viable method for the molecular-level study of minimal PKS enzymes from type II PKSs. Titers of 3 mg/L obtained from heterologous expression of *Kr*KSCLF in the *E. coli* BL21 cell line are a marked improvement over the heterologous expression of other KSCLFs in *S. coelicolor* CH999, which often afford less than1 mg/L of purified protein from a much more resource-intensive process.^23,29^

A workflow to gain access to high quantities of purified type II PKS enzymes opens the door to important molecular-level biochemical studies that can complement ongoing *in vivo* characterization of non-actinomycete type II PKS pathways. For example, recent pioneering work by the Takano Lab revealed that ancient orphaned type II PKS gene clusters from non-actinomycetes can be harnessed to synthesize type II polyketide upon reconstitution in *E. coli*.^13^ The *in vitro* biochemical studies of various ancient orphaned ACPs and KSCLFs, including *Kr*ACP and *Kr*KSCLF, complements this work by enabling access to the structural causes behind ACP/KSCLF compatibility. Information such as protein sequence, secondary structure, oligomeric state, and acylated state can be pooled together to create a catalog of ACPs and KSCLFs that in turn could be used to predict compatibility and design hybrid synthases capable of making novel chemical diversity.

While we were able to heterologously express and purify both components of the *K. racemifer* minimal PKS in *E. coli*, we did not observe any interaction between *Kr*ACP and *Kr*KSCLF *in vitro*. Although we do not currently know why these proteins are not interacting *in vitro*, it is possible that the size and charge of the amphipathic polyketide tunnel of *Kr*KSCLF (as compared to the actKSCLF)^23^ might play a role. The ability to obtain soluble *Kr*KSCLF enables further investigation of this possibility through site-directed mutagenesis experiments coupled with biophysical studies. Moreover, the role of acylation state in facilitating KSCLF – ACP interactions can be studied through pairing the chemoenzymatic acylation of the KSCLF and/or ACP with biophysical studies using previously established methods.^25,26^

Notwithstanding the lack of observed *Kr*ACP/*Kr*KSCLF interactions, the results presented herein lay the foundation for further investigation of *Kr*ACP and *Kr*KSCLF as potential components of a hybrid PKS and the methods described herein can be applied to explore other non-actinomycete type II PKS enzymes. The ability of *Kr*ACP to self-malonylate may represent an advantage of calling upon this ACP in chemical engineering efforts as it could participate in biosynthesis in the absence of a native malonyl transferase. Future ACPs heterologously expressed from non-actinomyces strains can be similarly tested for self-malonylation activity, and sequence alignments between ACPs with and without the ability to self-malonylate can potentially elucidate certain amino acid motifs that allow for ACP self-malonylation.

Taken together, our results support that non-actinomycete type II PKS enzymes can be robustly expressed and purified in *E. coli*. Access to these purified enzymes can be leveraged to increase knowledge of the structural determinants behind ACP/KSCLF compatibility. A large catalog of such information can enable more accurate prediction of compatibility between not just characterized but also the numerous and growing uncharacterized type II PKS ACPs and KSCLFs that have been identified through sequence analyses.

## Materials and Methods

### Construction of expression plasmids

The expression plasmid for the *E. coli* FAS AT FabD was provided as a gift from the Campopiano Research Group at University of Edinburgh. *Ktedonobacter racemifer* genomic DNA was purchased from the DSMZ-German Collection of Microorganisms and Cell Cultures (DSM 44963). Plasmids pYW2 and pkrACP (encoding the expression of *Ktedonobacter racemifer* KSCLF and ACP, respectively) were constructed by inserting amplified genes encoding the relevant enzymes into the NdeI/EcoRI restriction cut sites of pET28a vector via Gibson Assembly for pYW2 and restriction digestion ligation for p*Kr*ACP. See Supporting Information for the protein sequences of FabD, *Kr*ACP and *Kr*KSCLF.

The gene sequence encoding the expression of *Kr*KSCLF was amplified from the gDNA using the following primers (NdeI and EcoRI endonuclease cut sites are underlined): 5’CCTGGTGCCGCGCGGCAGC<u>CATATG</u>CGCCGTGTCGTTATCTCTG 3’ and 5’AGCTTGTCGACGGAGCTC<u>GAATTC</u>CTAGGCCCACGTGCGCAGT 3’. Thermal cycling conditions were as follows (40 ng DNA, Phusion polymerase): 1. 98 °C for 1 min, 2. 98 °C for 10 sec, 3. 72 °C for 2 min, 4. Repeat steps 2-3 a total of 34 times, 5. 72 °C for 10 min, 6. 4 °C forever. Amplified DNA was purified using the Zymoclean Recovery Kit (Zymo Research) and cloned into the NdeI and EcoRI endonuclease cut sides of linearized pET28a vector DNA via Gibson Assembly as follows. The insert and vector DNA were combined in a 3:1 insert:vector molar ratio in a volume of 5 μL and added 15 μL of Gibson Assembly Master Mix (New England BioLabs), consisting of T5 Exonuclease, Phusion Polymerase, Taq Ligase, dNTPs and MgCl2 in Tris-HCl buffer. The resulting mixture was incubated for 1 hr at 50 °C and transformed into DH5alpha and plated on LB agar supplemented with kanamycin (50 μg/mL). Plasmid DNA was isolated from colonies using a Qiagen QIAprep Spin Miniprep Kit and sequenced using standard T7 primers (Eurofins Genomics).

The gene sequence encoding the expression of *kr*ACP was amplified from the gDNA using the following primers (NdeI and EcoRI endonuclease cut sites are underlined): 5’AAAAAA<u>CATATG</u>GCTAAAGATTCGGG 3’ and 5’AAAAAA<u>GAATTC</u>TCATACGCTCTGGAC 3’.

Thermal cycling conditions were as follows (40 ng DNA, Phusion polymerase): 1. 98 °C for 1 min, 2. 98 °C for 10 sec, 3. 65 °C for 30 sec, 4. 72 °C for 30 sec, 4. Repeat steps 2-4 for a total of 34 times, 5. 72 °C 10 for minutes, 6. 4 °C forever. Amplified DNA was purified using the Zymoclean Recovery Kit (Zymo Research). Vector DNA was obtained by digesting pET28a vector with NdeI and EcoRI endonuclease enzymes, treating linearized vector with calf intestinal phosphatase (New England Biolabs) at 16 °C for 30 minutes, and subsequently purified with the Zymo Resaerch DNA Clean and Concentrator Kit (Zymo Research). Vector and insert were mixed in a 1:1 molar ratio with T4 ligase (0.25 μL; Thermo Fisher), 10X T4 ligase buffer (2 μL) and ddH_2_O to a total volume of 20 μL. After 3 hrs of incubating at 16 °C, the ligase was heat deactivated for 10 minutes at 65 °C, transformed into DH5alpha chemically competent cells, and plated on LB agar supplemented with kanamycin (50 μg/mL). Plasmid DNA was isolated from colonies using a Qiagen QIAprep Spin Miniprep Kit and sequenced using standard T7 primers (Eurofins Genomics).

### Expression and purification of krACP, krKSCLF, and FabD

Expression plasmids were transformed into chemically competent BAP1 cells^19^ (p*Kr*ACP) or BL21 cells (pYW2 (*Kr*KSCLF), pFabD (*E. coli* FAS AT) for expression. Seed cultures (10 mL LB, 50 μg kanamycin/mL) were grown at 37 °C with shaking and then added to production cultures (1 L LB, 50 μg kanamycin/mL). Production cultures were grown at 37 °C with shaking until OD600 was between 0.4 – 0.6. Each 1L culture was induced with 250 μL of 1 M Isopropyl β-D-1-thiogalactopyranoside (IPTG) and incubated at 18 °C for 18 – 21 h with shaking. Cells were harvested by centrifugation (5000 RPM, 4 °C, 20 min), resuspended in lysis buffer (50 mM sodium phosphate, 300 mM sodium chloride, 10 mM imidazole, pH 7.6) and sonicated on ice using an XL-200 Microson sonicator (40% A, 30 sec cycles, 10 min). Supernatant was clarified by centrifugation (13000 RPM, 1 hr), and the supernatant was incubated overnight on a nutating mixer at 4 °C with 2 mL nickel-NTA agarose slurry equilibrated in lysis buffer. After collecting the flow through, resin was washed with 75 mL of lysis buffer (50 mM sodium phosphate, 300 mM sodium chloride, 10 mM imidazole, pH 7.6), 100 mL of wash buffer (50 mM sodium phosphate, 300 mM sodium chloride, 30 mM imidazole, pH 7.6), and protein was eluted from the resin with 10 mL of elution buffer (50 mM sodium phosphate, 100 mM sodium chloride, 300 mM imidazole, pH 7.6). For KrKSCLF purification, 1 mM DTT and 10% glycerol were added to lysis, wash, and elution buffers. Proteins were concentrated using 3 kDa (*Kr*ACP, FabD) and 30 kDa (*Kr*KSCLF) Amicon Ultra (Millipore, 15 mL capacity) molecular weight cutoff (MWCO) centricons (4000 RPM, 4 °C) to A280 readings of 0.8-1.2. Prior to flash freezing, *Kr*ACP and FabD were dialyzed into 50 mM sodium phosphate, pH 7.6 using Thermo Fisher Slide-A-Lyzer dialysis cassettes. For SV-AUC and SEC studies, the *Kr*KSCLF sample was dialyzed into reducing buffer (50 mM sodium phosphate, 150 mM sodium chloride, 2 mM EDTA, 2 mM DTT, pH 7.2). Glycerol (10 % v/v) was added to the KrKSCLF samples. All protein aliquots were flash-frozen in liquid nitrogen and stored at −80 °C.

### *Kr*KSCLF purification by gel filtration chromatography

A GE Healthcare Superdex 75 increase 10/300 GL size exclusion column was equilibrated with 50 mM sodium phosphate, pH 7.6, 150 mM NaCl and 2 mM DTT. Next, the *Kr*KSCLF sample (600 μL, 49 μM) was injected using an ÄKTA Pure 25 L FPLC system. Flow rate was held at 0.8 mL/min and absorbance measured at 280 nm. Fractions (1 mL) were collected and those with a strong absorbance at 280 nm (fractions 4-6) were collected and concentrated by centrifugation for SV-AUC analysis.

### SDS-PAGE gel electrophoresis

Bio-Rad™ Mini-PROTEAN® TGX™ 10-well 30 μL 4-20 % precast polyacrylamide gels were used for SDS-PAGE gel electrophoresis. *Kr*KSCLF was dialyzed and diluted to 2x running concentration in reducing buffer (50 mM sodium phosphate, 150 mM sodium chloride, 2 mM EDTA, 2 mM DTT, pH 7.2). *Kr*ACP was dialyzed and diluted to 2x running concentration in 50 mM sodium phosphate, pH 7.6. Protein samples were further diluted 1:1 v/v to running concentration with 2x SDS-PAGE loading dye (100mM Tris-HCl, 4% w/v sodium dodecyl sulfate, 0.2% w/v bromophenol blue, 20% v/v glycerol, pH 6.8) to reach a final concentration of 1x SDS-PAGE loading dye. For *Kr*ACP samples run under reducing conditions, 5% v/v BME was added to the protein sample. Gels were run in tris-glycine running buffer (25 mM Tris, 190 mM glycine, 0.1% SDS, pH 8.3) at 120 V for 90 min. Gels were washed with ddH_2_O and stained for 15 minutes in Thermo Scientific™ GelCode Blue Safe Protein Stain (Coomassie G-250). After staining, gels were de-stained in ddH_2_O overnight or until sufficient band contrast had been reached prior to imaging.

### Tandem proteolysis mass spectrometry

Liquid chromatography – tandem mass spectrometry (LC-MS/MS) was performed by the Proteomics and Metabolomics Facility at the Wistar Institute (Philadelphia, PA). A Q Exactive Plus mass spectrometer (ThermoFisher Scientific) coupled with a Nano-ACQUITY UPLC system (Waters) were used for the analyses. Samples were digested in gel with trypsin and injected onto a UPLC Symmetry trap column (180 μm i.d. x 2 cm packed with 5 μm C18 resin; Waters). Tryptic peptides were separated by reversed phase HPLC on a BEH C18 nanocapillary analytical column (75 μm i.d. x 25 cm, 1.7 μm particle size; Waters) using a 95 min gradient formed by solvent A (0.1 % formic water in water) and solvent B (0.1 % formic in acetonitrile). A 30-min blank gradient was run between sample injections to minimize carryover. Eluted peptides were analyzed by the mass spectrometer set to repetitively scan *m/z* from 400 to 2000 in positive ion mode. The full MS scan was collected at 70,000 resolution followed by data-dependent MS/MS scans at 175,000 resolution on the 20 most abundant ions exceeding a minimum threshold 20,000. Peptide match was set as “preferred, exclude isotopes” option and charge-state screening were enabled to reject singly and unassigned charged ions. Peptide sequences were identified using the MaxQuant 1.6.1.0 program. MS/MS spectra were searched against a custom *E. coli* UniProt protein database using full tryptic specificity with up to two missing cleavages, static carboxamidomethylation (57.02146) of Cys and variable oxidation (15.99491) of Met. Consensus identification lists were generated with false discovery rates of 1% at protein, peptide and site levels. Data were plotted by the Wistar Institute.

### High-performance liquid chromatography (HPLC)

HPLC data was collected using a Varian ProStar High-PerformanceLiquid Chromatography instrument equipped with a Macherey-Nagel NUCLEOSIL^®^ C4 column. The instrument was equipped with two solvents: Solvent A (H_2_O + 0.1% trifluoroacetic acid) and Solvent B (acetonitrile + 0.1% trifluoroacetic acid). The column was equilibrated for 10 minutes at a 1 mL/min flow rate with 90% Solvent A and 10% Solvent B, after which 25 μL of 50 μM krACP was injected into the column. The column was then run for 30 minutes, during which the solvent gradient linearly increased from 10% Solvent B to 100% Solvent B (3% min^−1^). The eluent was measured by absorbance at 235 nm. Data was plotted in Microsoft Excel (v.15.28)

### Liquid chromatography-mass spectrometry and phosphopantetheine ejection assay

LCMS data was collected in the positive mode on an Agilent Technologies InfinityLab G6125B LC/MSD coupled with Agilent 1260 Infinity II LC system with a Waters XBridge Protein BEH C4 reverse phase column. The following solvent gradient was used (solvent A = water + 0.1% formic acid; solvent B = acetonitrile + 0.1% formic acid): 0–1 min 95% A, 3.1 min 5% A, 4.52 min 5% A, 4.92–9 min 95% A. Samples were run using a capillary voltage of 3000 V and a fragmentation voltage of 75V. For the Ppant ejection assay, the fragmentation voltage was adjusted to 250V. Resulting spectra were deconvoluted using ESIProt.^30^ Data was plotted in Microsoft Excel (v.15.28)

### *Kr*ACP malonylation experiments

To a microcentrifuge tube, *Kr*ACP (2.6 mM, 26.9 μL) was diluted to a volume of 500 μL in 50 mM sodium phosphate, pH 7.6 to reach a target concentration of 140 μM *Kr*ACP. To a second tube, *Kr*ACP (2.6 mM, 26.9 μL) and malonyl-CoA (50 mM, 10 μL) was added and diluted to a target volume of 500 μL in sodium phosphate, pH 7.6 to reach target concentrations of 140 μM *Kr*ACP and 1 mM malonyl-CoA. To a third tube, *Kr*ACP (2.6 mM, 26.9 μL), malonyl-CoA (50 mM, 10 μL), and FabD (56.1 μM, 13.4 μL) were mixed in 50 mM sodium phosphate, pH 7.6 to reach target concentrations of 140 μM *Kr*ACP, 1 mM malonyl-CoA, and 1.5 μM FabD in a total volume of 500 μL. All solutions were incubated at 23 °C for 30 min.

The 500 μL reaction mixtures were loaded onto a GE Healthcare Superdex 75 increase 10/300 GL size exclusion column attached to a ÄKTA Pure 25 L FPLC system. Samples were run at a flow rate of 0.8 mL/min and absorbance was measured at 280 nm. Fractions were collected at 1 mL, and Fraction 8 (corresponding to 12.7-13.7 mL volume eluted) was collected for the Ppant ejection assay analysis. 25μL samples of the collected fractions were run on LCMS using a capillary voltage of 3000 V and a fragmentation voltage of 250 V. Data was plotted in Microsoft Excel (v.15.28)

### Sedimentation velocity analytical ultracentrifugation (SV-AUC)

Sedimentation velocity experiments were performed using a Beckman Optima XL-A ultracentrifuge equipped with a 4-hole An-60 Ti rotor. Temperature-corrected partial specific volumes, density and viscosity were calculated using Sednterp (beta version).^31^ Data were analyzed using the dc/dt method DCDT^+^ (v.2.4.0), and a model-independent continuous c(s) distribution from Sedfit (v. 15.3).^32^ Confidence intervals for temperature-corrected sedimentation coefficient, s(*20,w*), diffusion coefficient, D(*20,w*), and molecular mass (kDa) were computed using bootstrap method, CI 90% (± 1.65 sigma) implemented in DCDT^+^ (v.2.4.0).

SV-AUC experiments used *Kr*KSCLF purified by gel filtration chromatography in elution buffer (50 mM sodium phosphate, pH 7.6) with the presence (**Fig 6**) or absence of salt (150 mM NaCl) and reducing agent (2 mM DTT). An SV-AUC concentration gradient profile of *Kr*KSCLF prior to gel filtration chromatography purification was also taken (**Fig S4B**). For *Kr*KSCLF purified in the presence of salt and reducing agent, *Kr*KSCLF concentration was 2.9 μM and *Kr*ACP concentration was 140 μM. A mixed sample containing both *Kr*KSCLF (4 μM) and *Kr*ACP (110 μM) was also analyzed. All samples were diluted to the required starting absorbance between 0.3 and 1.0 AU and had a final concentration of 150 mM NaCl and 1 mM DTT. SV-AUC runs used two-channel Epon charcoal-filled centerpieces containing 450 μL protein samples and 450 μL buffer as reference. Sedimentation velocity boundaries were measured at a speed of 42,000 rpm at 20 °C using a step size of 0.003 cm, a delay time of 0 sec, and a total of 150 scans. Samples were monitored at 280 nm.

Sample heterogeneity was determined using model-independent continuous c(s) distribution where regularization of the distribution by the maximum entropy with the parameter α constrained value at 0.95. The s and D parameters required to solve the Lamm equation are calculated with the approximation that all species in a solution for a given sample have similar signal-average frictional coefficient f/f0. The distributions were plotted using the Gussi interface (v. 1.0.3) implemented in Sedfit (v. 15.3).

### Circular dichroism spectroscopy (CD)

CD spectra were collected using an Aviv Model 410A circular dichroism spectropolarimeter. Proteins were dialyzed in absence of DTT to prevent any light scattering from DTT. *Kr*KSCLF was dialyzed overnight into 50 mM sodium phosphate, pH 7.6, and *Kr*ACP was dialyzed overnight into 10 mM Tris-HCl, pH 7.6. Protein samples were diluted to 200 μL to final concentrations of 13 μM in 50 mM sodium phosphate, pH 7.6 for *Kr*KSCLF and 50 μM into 10 mM Tris-HCl, pH 7.6 for *Kr*ACP. Prior to injection, samples were filtered using a 0.2 μM low protein-binding filter with a HT Tuffryn Membrane (Pall Corporation). Samples were injected into a High Precision Quartz SUPRSIL cuvette with 0.1 cm pathlength (Hellma Analytics). The spectropolarimeter was purged with nitrogen for one hour, and CD spectra were collected at 25 °C with a range of 260-180 nm using a bandwidth of 1 nm, a 0.5 nm step size, and an averaging time of 3 sec. Changes in signal at 222 nm were followed as function of temperature using the following parameters; 10-90 °C, 2-min equilibration, heating rate 2 °C min^−1^, 30 sec signal averaging time, and 1 nm bandwidth. Pre- and post-Tmelt spectra were smoothed using a smoothing function implemented in the Aviv software, applying a window width of 11 data points, degree 2. Thermal stability data was analyzed using CDpal (v. 2.18) using a two-state unfolding model N↔D with the standard assumption that ΔCP = 0. Estimated errors for each fitted parameter were calculated using a Jackknifing method implemented in CDpal (v.2.18). Data was plotted in Origin (v.8.6.0). The resulting spectra were converted to units of mean residue ellipticity (MRE) using the amino acid sequence of the proteins, smoothed, and normalized to an MRE of 0 at 250 nm.

## Acknowledgements

We are grateful for generous support from the National Science Foundation CAREER Award (CHE-1652424 to L.K.C.) and the Henry Dreyfus Teacher-Scholar Award (TH-10-2020) to L.K.C. We would like to thank Grayson S. Hamrick and Sharon C. Nwankpa for their technical support with protein expression and purification, Professor Robert Fairman (Haverford Biology) for generously offering us the use of the analytical ultracentrifuge for this work, Professor Dominic Campopiano (University of Edinburgh) for providing the FabD expression plasmid, and Dr. Maureen Hillenmeyer (Hexagon Bio) for helpful discussions).

## Supporting Information for

### Protein sequences of expressed proteins

#### *Kr*ACP

MGSSHHHHHHSSGLVPRGSHMAKDSGQATVSSRVYEILRPYAEGIALTPETHLSDDLNIDSIELVEVGVALE KEFSNKRFTITGLKSCPTVQDLVKLVEQTVAAEQVQSV*

#### *Kr*KSCLF

MGSSHHHHHHSSGLVPRGSHMRRVVISGLGVVSPAGIGKEAFWQNLLDGVSGAVALDRVTCSPLFGHHEF GAQAVCEVTGFDPVTHHVPTAYHSADRFIQFAFAAVHQAFLDAHLDQDTWDVSRVGVTLATAICGTQTLDLEFTKSTNGGKEAFHSEGISPFLYSAAMGNSAALAVGSRYGLQGECATLSTGCIAGLDAISYAYESIAYGD HDVMIAGASEAPITPITIAAFDIINCLSHHSDPTTASRPFSVDRDGFVLSEGCGIVVLEELEHALARNAPIYAEILGCDVTEHAVHMTDMSPEGRDLARAITGALGKANVEPEAIDFVNAHGTSTPQNDFFESLALKTSLGQQALTIPVNSTKSMVGHALAAASAVEVVACAMSMQTQRIHPTINLVNPDPRCDLDYVPNHARSHDVQRMLTTASGFSGLHAAMVLESYSTREEMMARIVITGMGVISPYGIGSGILWEKLLAGENGLKPLTTFETSHIQCRVGGQLLDFRPEAYLSPRLIRKIDRFSTFGLISAYLALQDAGLLSDGKKPVWTQQEQHSHRVGITVGNNLGGWEFAERELRHLWALGPRDVSPHMATAWFPAAVQGNMSIYFGIKGIGRTFLSDRASGALAIMHAADCLQRGRADIMLAGGTEAPFSPYAALCYETSGLMSKKAVTGSPETYRPFDEAHDGLVAGEGAAFFILERAEDAEKRGATILAEVAGWASTNDGYHPVQPAPDGERYAAAMTRAMQRADASADEVDCLFAAGSAVPDEDVSETRAIHLALREAVRRVPVATPKSAFGNLFGAAFPVDMAIALLAMQHRVLPATLHLDQAAPGCDLDYVPQTPRSVDHLDRCLINARGIGGANASVLLRTWA

#### *E. coli* FabD

MGSSHHHHHHSSGLVPRGSHMTQFAFVFPGQGSQTVGMLADMAASYPIVEETFAEASAALGYDLWALTQQGPAEELNKTWQTQPALLTASVALYRVWQQQGGKAPAMMAGHSLGEYSALVCAGVIDFADAVRLVEMRGKFMQEAVPEGTGAMAAIIGLDDASIAKACEEAAEGQVVSPVNFNSPGQVVIAGHKEAVERAGAACKAAGAKRALPLPVSVPSHCALMKPAADKLAVELAKITFNAPTVPVVNNVDVKCETNGDAIRDALVRQLYNPVQWTKSVEYMAAQGVEHLYEVGPGKVLTGLTKRIVDTLTASALNEPSAMAAALEL

### Protein threading and modeling

Plasmid nucleotide sequences were translated using the web program ExPasy. This nucleotide sequence was threaded by the web program Phyre 2.0. Modeling and analysis of the threaded sequence was performed using PyMol.

**Fig S1.**
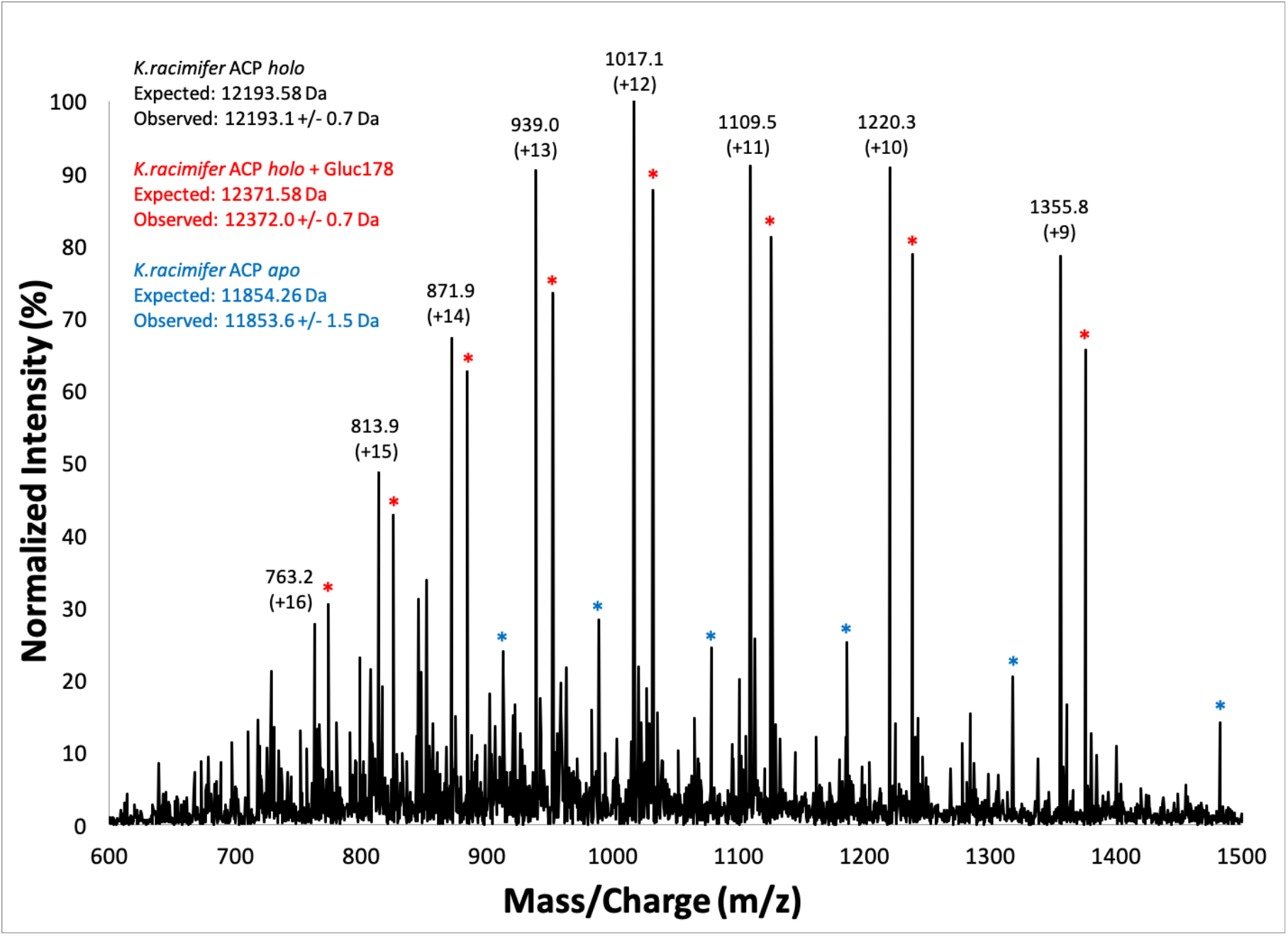
LC-MS spectrum of purified krACP. The *Kr*ACP LS-MS spectrum shows both *holo* and *apo*-*Kr*ACP post-purification in the *E. coli* BAP1 competent cell line. All theoretical molecular weights are calculated with the removal of a single methionine residue.

**Fig S2.**
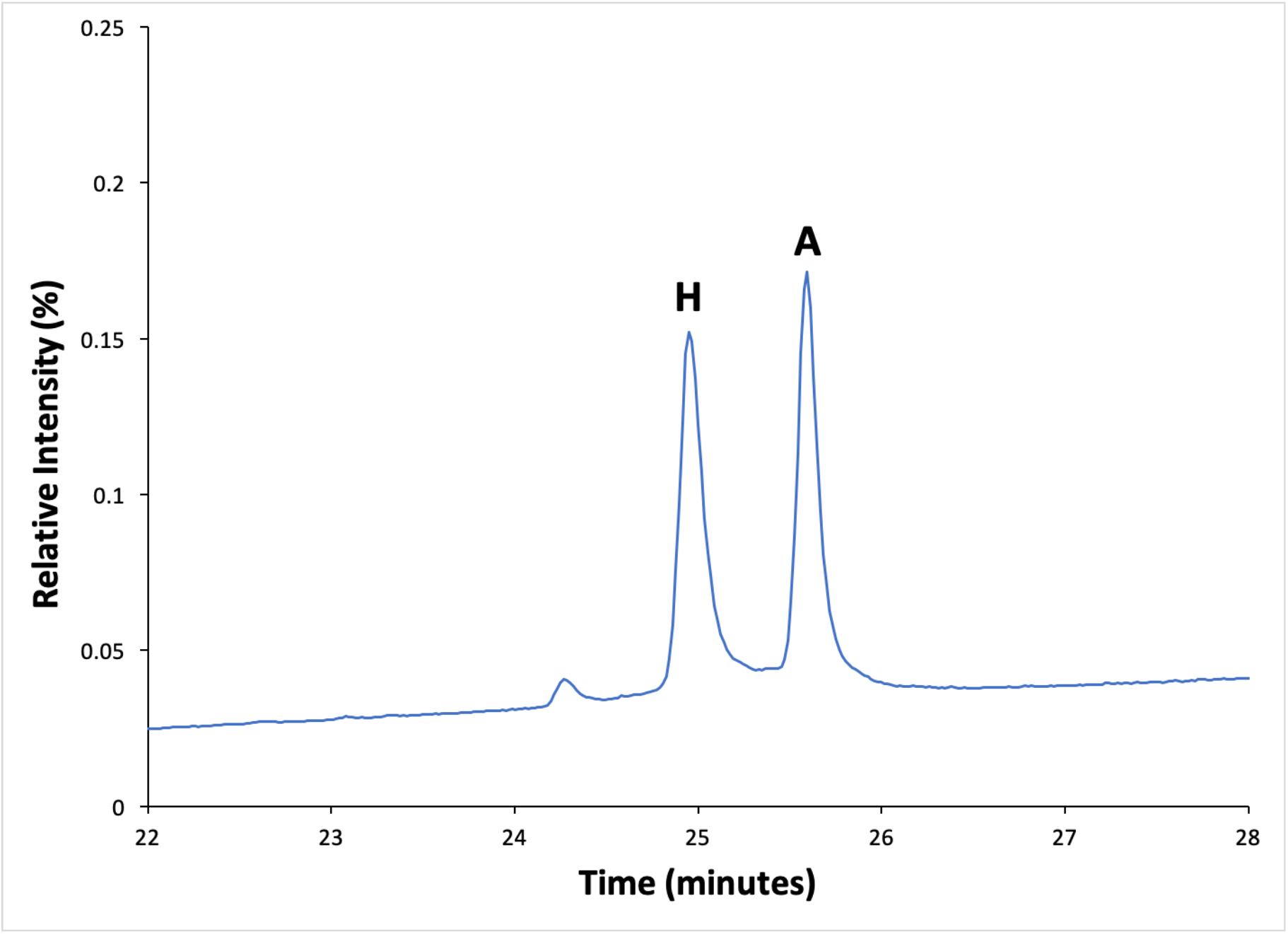
High-pressure liquid chromatography profile of *Kr*ACP. H: Peak corresponding to *holo*-*Kr*ACP. A: Peak corresponding to *apo*-*Kr*ACP. The HPLC profile of purified *Kr*ACP confirms the presence of both *holo* and *apo*-*Kr*ACP in an approximate 1:1 ratio.

**Fig S3.**
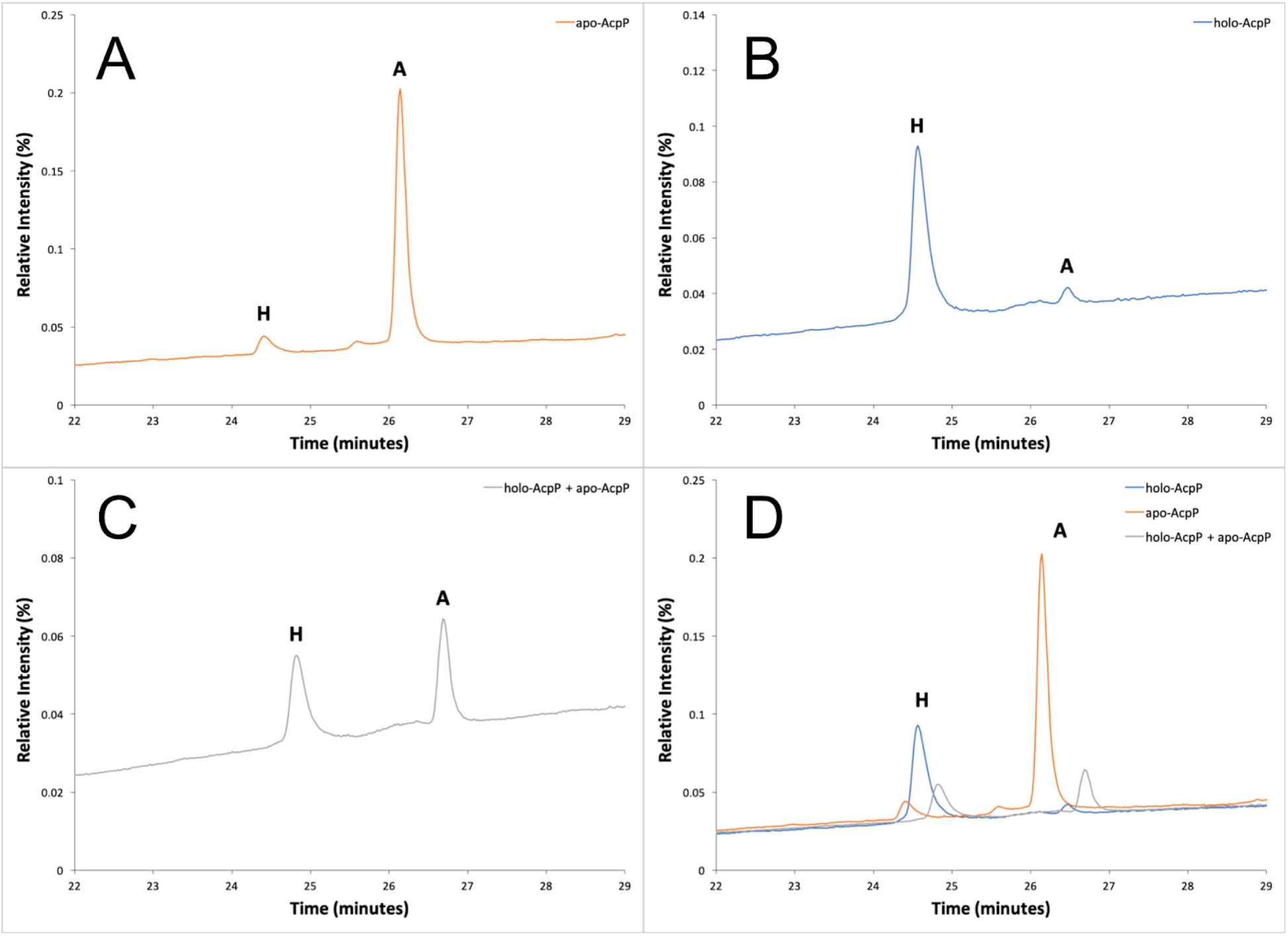
High-pressure liquid chromatography spiking experiments with *E. coli* AcpP. H corresponds to *holo*-AcpP peak and A corresponds to *apo*-AcpP peak. A. HPLC profile of a primarily *apo*-AcpP sample (70 μM) shows a minor peak at 24.41 minutes corresponding to *holo*-AcpP and a major peak at 26.14 minutes corresponding to *apo*-AcpP; B. HPLC profile of a primarily *holo*-AcpP sample (70 μM) shows a major peak at 24.59 minutes corresponding to *holo*-AcpP and a minor peak at 26.47 minutes corresponding to *apo*-AcpP; C. HPLC profile of a mixed sample containing (52.5 μM) *holo*-AcpP and (17.5 μM) *apo*-AcpP shows major peaks at 24.80 minutes corresponding to *holo*-AcpP and at 26.62 minutes corresponding to *apo*-AcpP; D. Overlaid HPLC profiles of subfigures S3 A-C.

**Fig S4.**
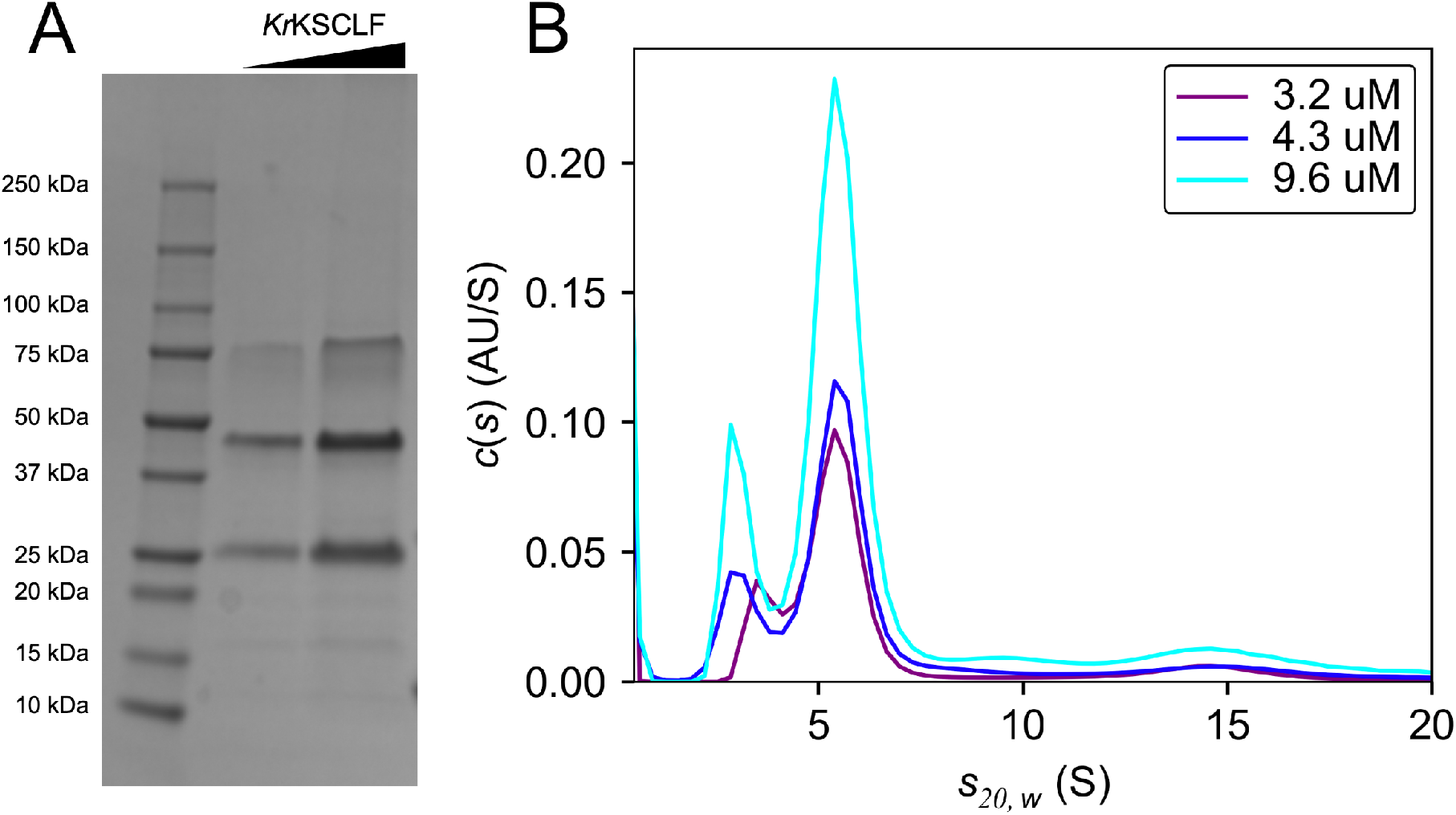
Pre-SEC SDS-PAGE and SV-AUC analysis of *Kr*KSCLF. A. SDS-PAGE of *Kr*KSCLF pre-SEC purification. Lanes: 1. Protein standards ladder; 2. *Kr*KSCLF (10 μL, 3.7 μM); 3. *Kr*KSCLF (20 μL, 3.7 μM). B. SV-AUC profile of *Kr*KSCLF pre-SEC purification at increasing concentrations. The distribution between the 25.0 kDa peak and the 98.7 kDa peak of SEC-purified *Kr*KSCLF does not change with increasing *Kr*KSCLF concentration. This indicates that the protein(s) represented by the 25.0 kDa peak is/are not in equilibrium with the protein(s) presented by the 98.7 kDa peak, and that the two peaks do not represent different oligomeric states of the same protein. This 25.0 kDa peak persists after SEC purification in 50 mM sodium phosphate, pH 7.6, but can be separated upon addition of 150 mM NaCl and 2 mM DTT to the equilibration buffer.

**Fig S5.**
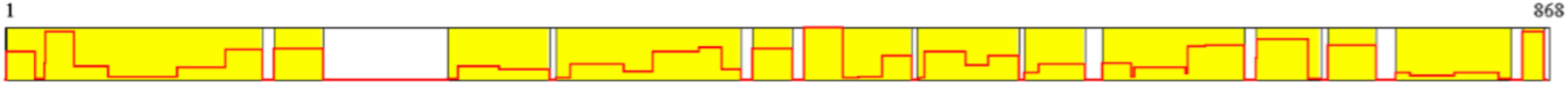
Tandem proteolysis mass spectrometry profile of ~47 kDa SDS-PAGE Band. The ~47 kDa SDS-PAGE band was digested with proteolytic enzymes and the resulting mass spectra fragments was compared to the mass of expected proteolysis cut sites. Comparing the tandem proteolysis mass spectrometry profile of the ~47 kDa band to expected proteolysis cut sites in the full *Kr*KSCLF sequence, all expected proteolysis fragments from both the *Kr*KS and *Kr*CLF sequences are present in the band. Blank fragments were either too small or too large to detect via this method.

**Fig S6.**
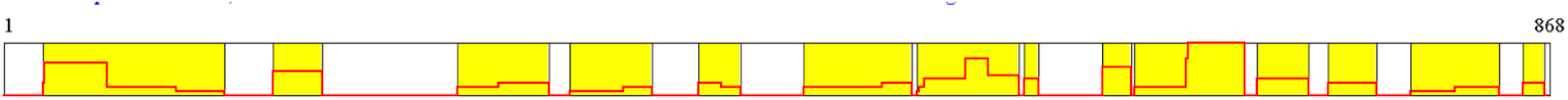
Tandem proteolysis mass spectrometry profile of ~25 kDa SDS-PAGE band. The ~25 kDa SDS-PAGE band was digested with proteolytic enzymes and the resulting mass spectra fragments was compared to the mass of expected proteolysis cut sites. Comparing tandem proteolysis mass spectrometry profile of the ~25 kDa band to expected proteolysis cut sites in the full *Kr*KSCLF sequence, most expected proteolysis fragments from both *Kr*KS and *Kr*CLF are present in the band. However, the band was mostly comprised of the *E. coli* proteins SlyD, ThiM, and TpiA.

**Fig S7.**
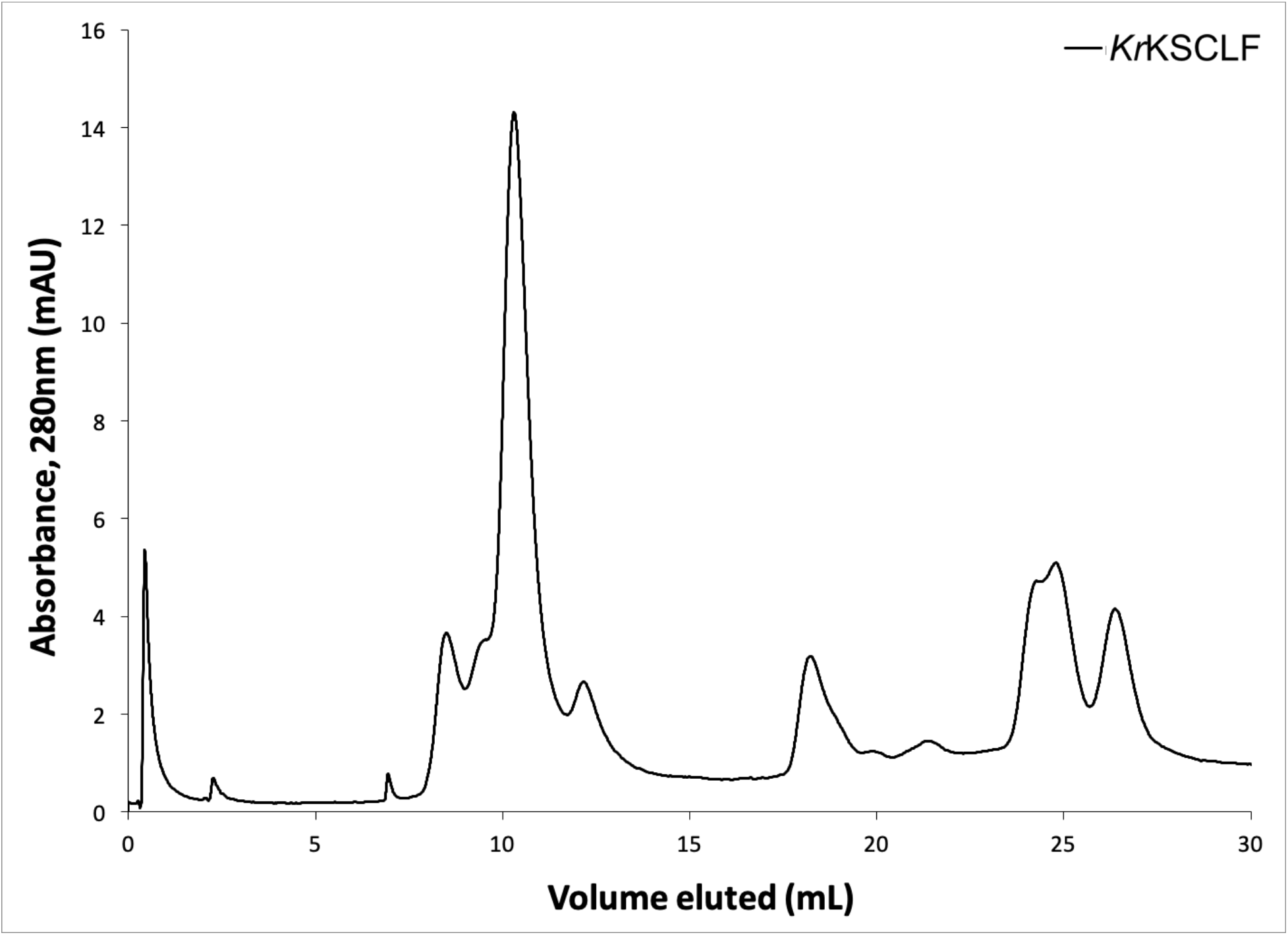
SEC purification of *Kr*KSCLF. The SEC profile of *Kr*KSCLF shows a major species at a high molecular weight, attributed to the *Kr*KSCLF heterodimer. This peak (Fractions 4+5, corresponding to 8.7-10.7 mL eluted) was collected and further analyzed by SDS-PAGE and SV-AUC.

**Fig S8.**
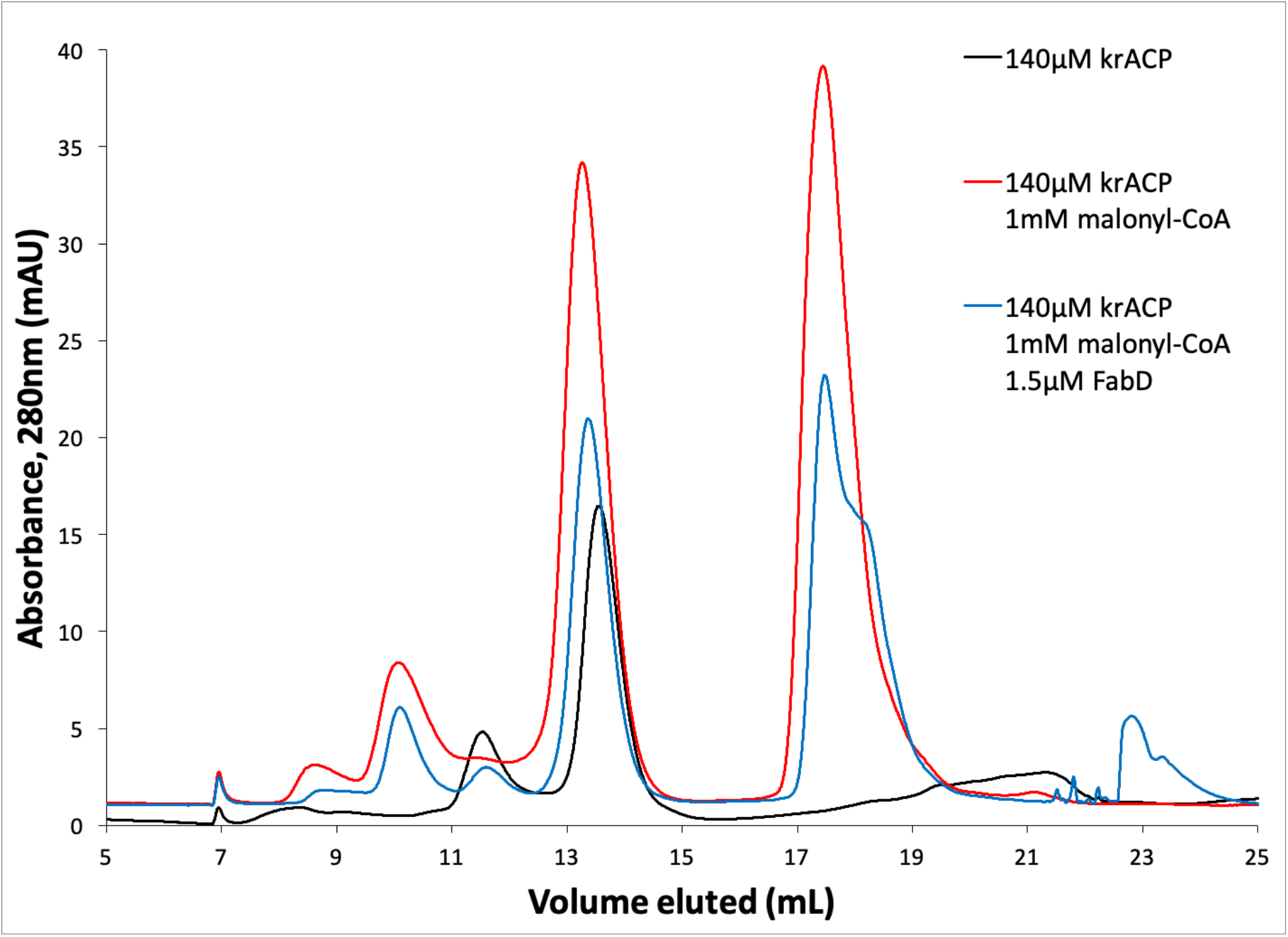
SEC Purification of *Kr*ACP malonylation solutions. The fraction corresponding to 12.7-13.7 mL volume eluted was collected and analyzed by LC-MS.

**Fig S9.**
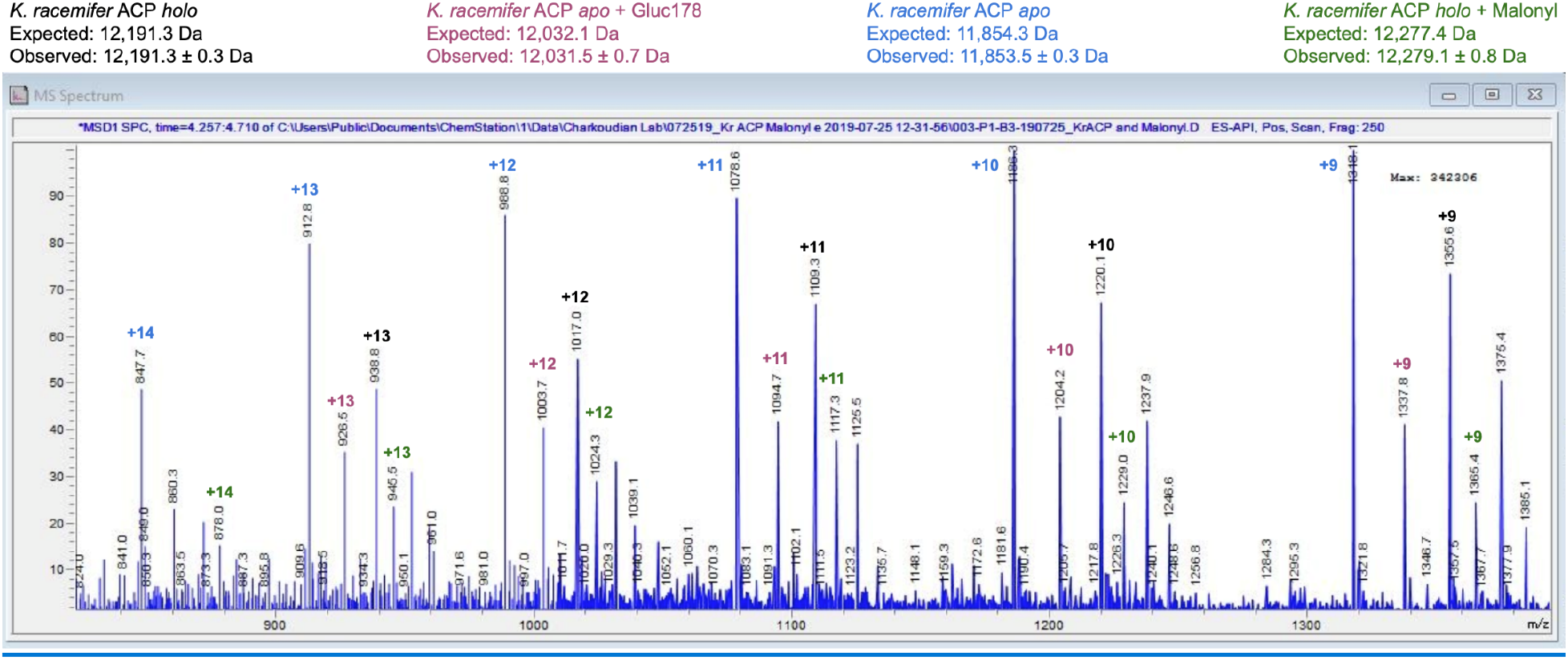
LC-MS spectrum of *Kr*ACP after incubation with malonyl CoA. The *Kr*ACP LC-MS spectrum shows the presence of *apo*-(red, blue), *holo-* (black), and *malonyl-* (green) *Kr*ACP after incubation with malonyl CoA. All theoretical molecular weights are calculated with the removal of a single methionine residue.

**Figure S10.**
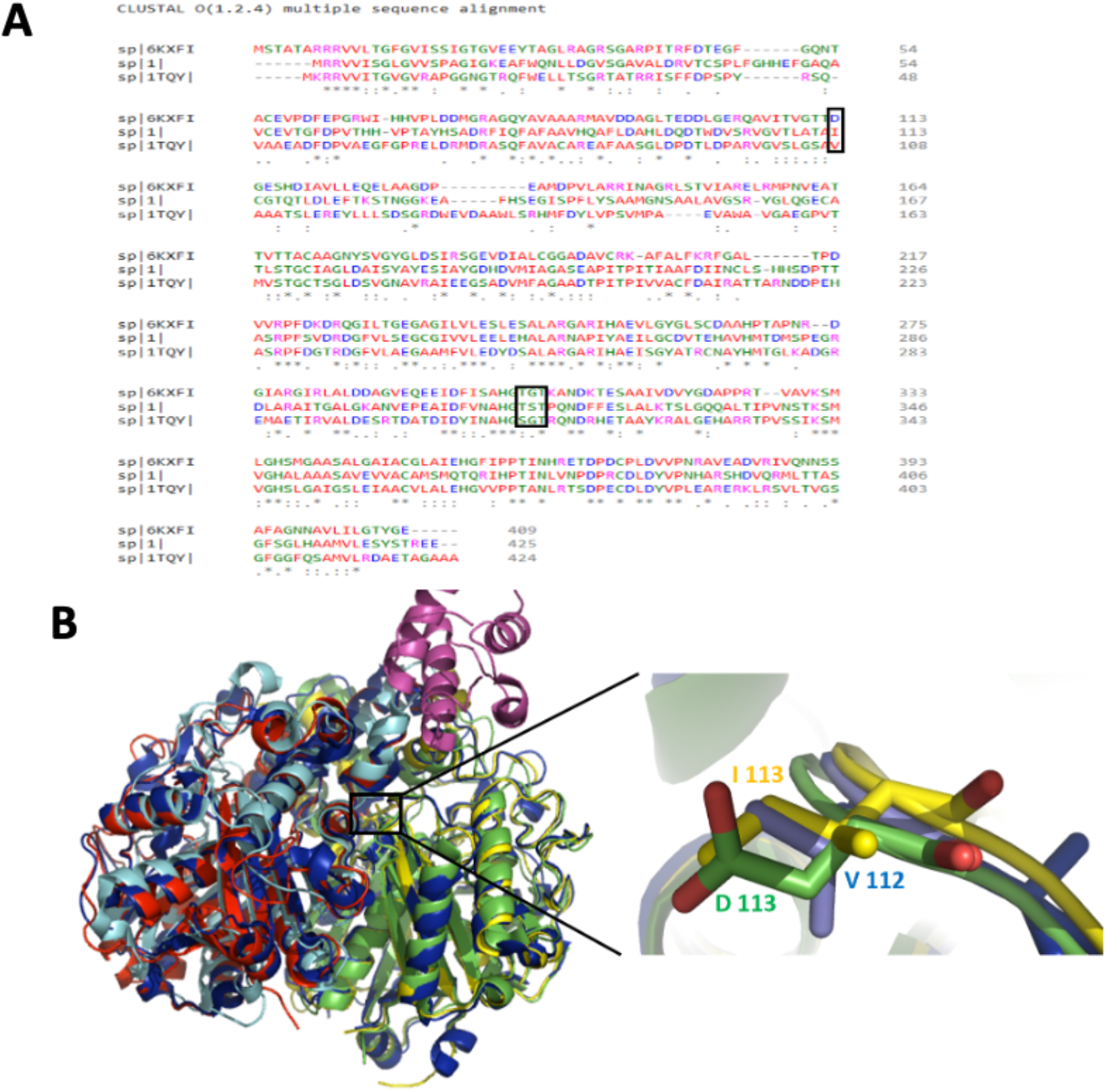
Sequence (A) and structure (B) alignments of *Kr*KS, actKS (1TQY) and Ig11 (6KXFI). Green (Iga11), yellow (KrKS), blue (actKS).

**Fig S11.**
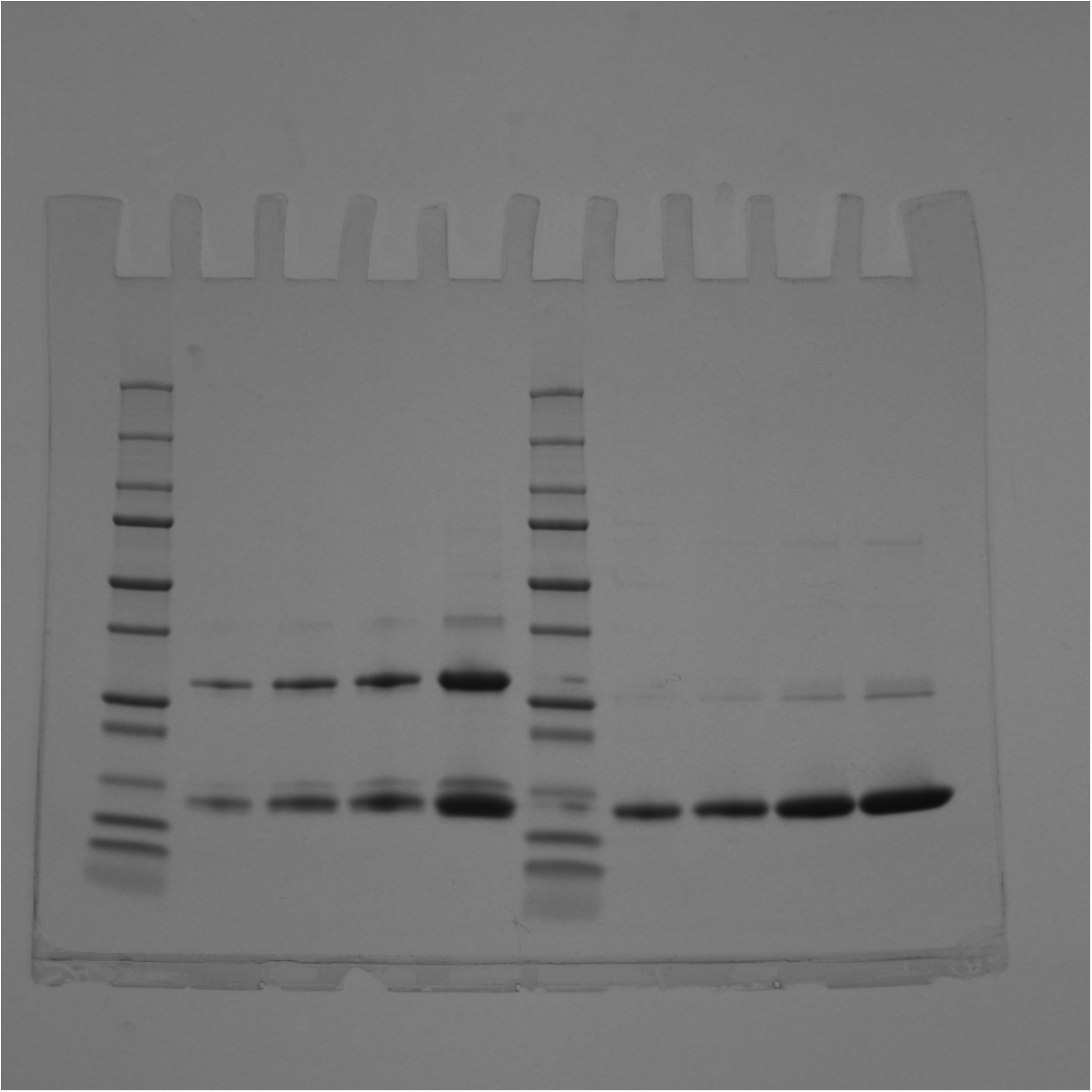
Uncropped Gel of Fig 2A. *Kr*ACP concentration gradients run both without and with the presence of 5% v/v BME. Lanes: 1. Protein standards ladder; 2. *Kr*ACP (31.4 μM, 5 μL) run under non-reducing conditions; 3. *Kr*ACP (31.4 μM, 10 μL) run under non-reducing conditions; 4. *Kr*ACP (31.4 μM, 15 μL) run under non-reducing conditions; 5. *Kr*ACP (31.4 μM, 20 μL) run under non-reducing conditions; 6. Protein standards ladder; 7. *Kr*ACP (31.4 μM, 5 μL) run under reducing conditions; 8. *Kr*ACP (31.4 μM, 10 μL) run under reducing conditions; 9. *Kr*ACP (31.4 μM, 15 μL) run under reducing conditions; 10. *Kr*ACP (31.4 μM, 20 μL) run under reducing conditions.

**Fig S12.**
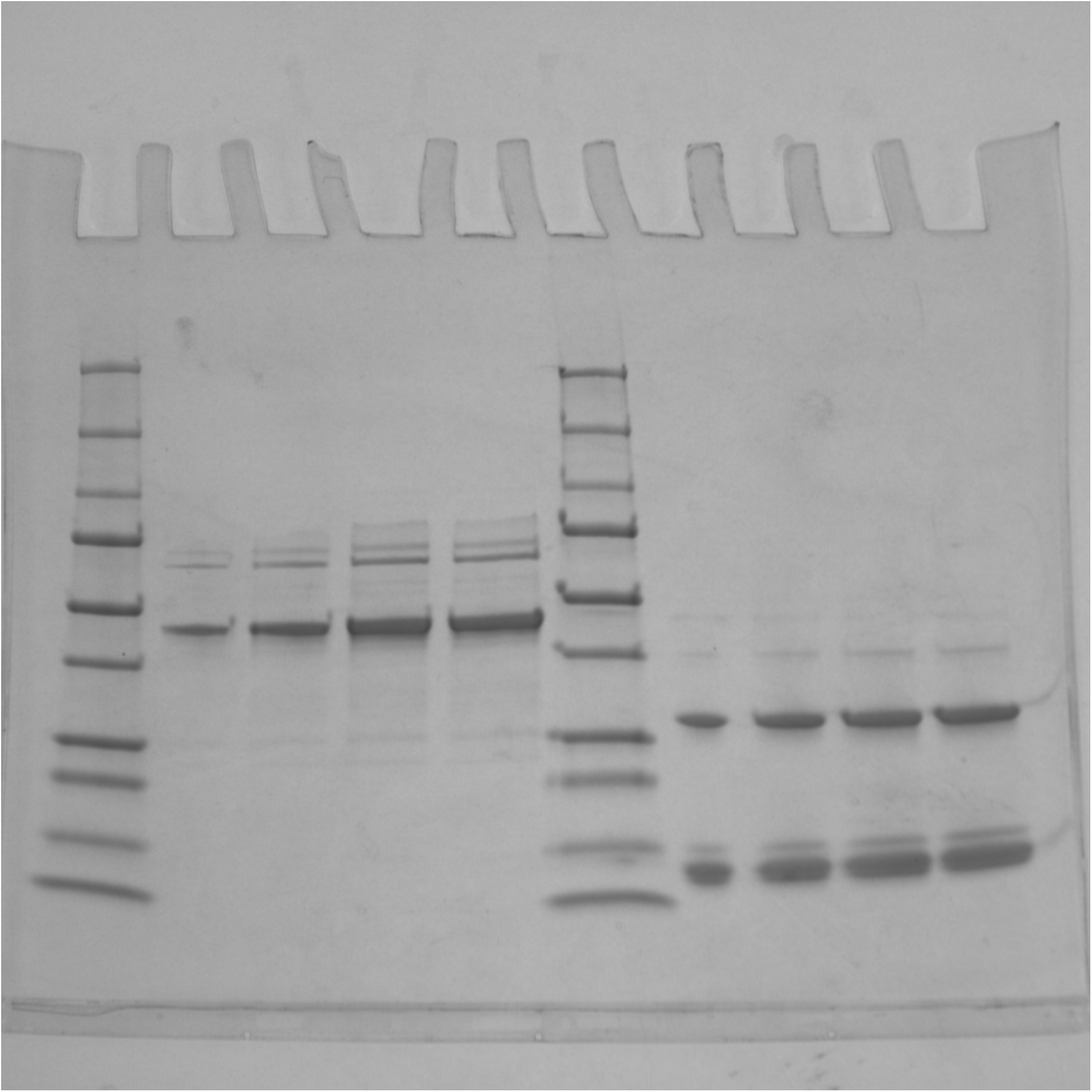
Uncropped Gel of Fig 2B. *Kr*KSCLF purified under reducing conditions. 1. Protein standards ladder; 2. *Kr*KSCLF (5 μL, 2.6 μM); 3. *Kr*KSCLF (10 μL, 2.6 μM); 4. *Kr*KSCLF (15 μL, 2.6 μM); 5. *Kr*KSCLF (20 μL, 2.6 μM); 6. Protein standards ladder; 7. *Kr*ACP (5 μL, 25 μM); 8. *Kr*ACP (10 μL, 25 μM); 9. *Kr*ACP (15 μL, 25 μM); 10. *Kr*ACP (20 μL, 25 μM).

**Fig S13.**
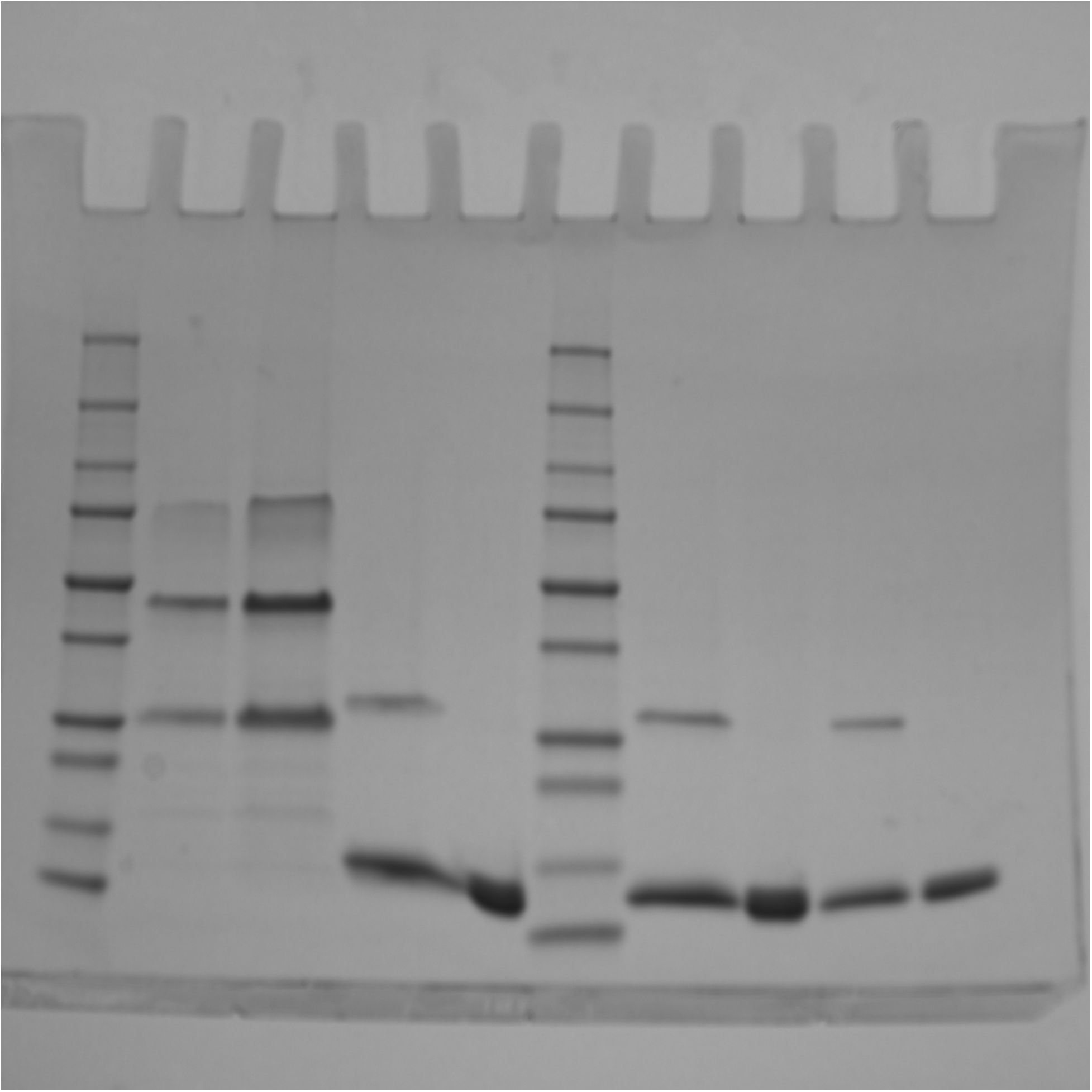
Uncropped Gel of S4A Fig. *Kr*KSCLF purified under reducing conditions pre-SEC purification. 1. Protein standards ladder; 2. *Kr*KSCLF (10 μL, 3.7 μM); 3. *Kr*KSCLF (20 μL, 3.7 μM); 4. *Kr*ACP (15 μL, 25 μM); 5. *Kr*ACP + 5% v/v BME (15 μL, 25 μM); 6. Protein standards ladder; 7. *Kr*ACP (10 μL, 25 μM); 8. *Kr*ACP + 5% v/v BME (10 μL, 25 μM); 9. *Kr*ACP (5 μL, 25 μM); 10. *Kr*ACP + 5% v/v BME (5 μL, 25 μM).

